# Natural variation in the *Arabidopsis thaliana* root and shoot response to boron deficiency reveals sensitive and responsive phenes and phylogenetic and geographic clustering of boron efficiency adaptations

**DOI:** 10.1101/2024.07.08.602492

**Authors:** Thomas D. Alcock, Manuela Désirée Bienert, Astrid Junker, Henning Tschiersch, Rhonda C. Meyer, Sreelekha Kudamala, Nicolaus von Wirén, Thomas Altmann, Gerd Patrick Bienert

## Abstract

- Boron (B) is an essential plant micronutrient with a crucial role in cell wall stability. To improve plant tolerance to suboptimal B-availability, it is crucial to understand the mechanisms plants have evolved to tolerate B-limited conditions.
- We assessed temporal physiological, ionomic and molecular responses to B-deficiency across a diversity panel of 185 *Arabidopsis thaliana* accessions. Plants were characterised in soil-substrate in an automated phenotyping system to track shoot performance, and on agar plates to track root traits.
- Accessions varied widely in their response to B-deficiency. Rosette area decreased by 50% under B-deficiency across all accessions on average but by less than 15% in B-efficient accessions alone. Most root traits suffered under B-deficiency but lateral root length increased in 20% of accessions. The B-uptake and B transporter expression response of B-efficient and B-inefficient accessions was similar, suggesting increased B-use rather than B-uptake efficiency in efficient accessions. Phylogenetic analysis demonstrated enrichment of B-efficient accessions within a clade containing accessions originating from areas characterised as B-deficient. *K*-means clustering revealed increased lateral root length was associated with efficient shoot growth under B-deficiency.
- The results demonstrate different response strategies to B-deficiency, indicating target traits to improve the performance of plants grown under B-limiting conditions.

## Introduction

Boron (B) is an essential trace element for vascular plants (Lewis, 2020; Wimmer et al., 2020). Boron forms dimerizing di-ester bonds between two rhamnogalacturonan II (RG-II) monomers in the pectin fraction of the primary cell wall. The number of RG-II dimers determines the integrity, plasticity, elasticity and stability of the plant cell wall (O’Neill et al., 2004). Hence, at the cellular level, B is biochemically important for determining cell wall structure, apoplastic transport and compartmentalisation processes, stabilisation of membranes, cell growth, cell elongation, cell differentiation, pollen development, pollen tube growth, mechano-sensing, and penetration resistance to pathogens. At the macroscopic level, B-deficiency results in disease-like symptoms such as heart rot in roots and shoots of crops such as rapeseed, broccoli or sugar beet (Bergmann, 1992), susceptibility towards pathogens, impaired root and shoot meristem activity, morphological abnormalities such as stunting, leaf curling, wilting, and leaf darkening, and impaired flower development and fertility (Marschner, 2012). Together, these impairments lead to decreased plant performance and stress tolerance and ultimately reduced yield (Shorrocks, 1997; Eggert and von Wirén, 2013).

Worldwide, B-deficiencies frequently cause yield- and quality losses in numerous agricultural and horticultural crops. This is largely due to many areas used for agriculture containing insufficient soil B concentrations to meet the demands of diverse crop species throughout the whole growing season or during temporary periods of high B demand by the crop (Shorrocks, 1997). Even cereals, which have a relatively low demand for B (Bienert et al., 2023), can suffer from yield losses due to B-deficiency during the critical period of fertilisation. In addition, increasingly common climatic extremes can increase the occurrence and severity of B-deficiency-induced symptoms, particularly when periods of intense rainfall (resulting in B leaching) are followed by a persistent drought (preventing B transport towards the plant roots due to a restricted soil-water mass flow). At their onset and during their early stages, B-deficiency-induced growth defects are typically phenotypically latent, without macroscopic symptoms. Thus, reliable field-suitable biomarkers for the onset of B-deficiency in crops are lacking, and modern B fertilisation management strategies are not always employed when most needed. The breeding of B-efficient crop cultivars is crucial to sustainably improve the performance and fertility of crops grown under temporary suboptimal B conditions. However, to achieve this, it is necessary to understand physiological and molecular mechanisms underlying B-efficiency in plants.

Plant nutrient homeostasis and efficiency mechanisms have been identified based on studies addressing natural variation in the ionome of the model plant *Arabidopsis thaliana* (Rus et al., 2006; Loudet et al., 2007; Baxter et al., 2008, 2012; Kobayashi et al., 2008; Morrissey et al., 2009; Chao et al., 2012; Pineau et al., 2012; Koprivova et al., 2013, Campos et al., 2017). With respect to mechanisms regulating plant B status, employing forward and reverse genetic approaches with *A. thaliana* has led to the identification and functional and molecular characterisation of genes and proteins regulating B uptake and transport (Onuh and Miwa, 2021). In *A. thaliana*, two transmembrane protein families are chiefly responsible for this, namely the boron transport (BOR) and Nodulin26-like intrinsic protein (NIP) families. Members of the BOR transporter family form functional secondary active carrier proteins that mediate the efflux of B in form of borate out of the cytoplasm. The NIP family belongs to the aquaporin superfamily and NIP proteins are bidirectionally permeable to B in form of boric acid (Onuh and Miwa, 2021). The AtNIP5;1 protein was shown to be essential for the efficient uptake of B from the soil into the plant under B-deficient growth conditions (Takano et al., 2006). AtNIP6;1 is important for the preferential translocation of B out of the vasculature into young growing leaves (Tanaka et al., 2008). The tapetum-specific boric acid channel AtNIP7;1, was demonstrated to be involved in maintaining fertility under low B conditions (Routray et al., 2018). In addition, pollen-specific AtNIP4;1 and AtNIP4;2 have been shown to be important for pollen tube elongation under limited B conditions (Di Giorgio et al., 2016). AtBOR1 acts synergistically with AtNIP5;1 under low B supply and is required for loading B into the xylem in roots, and therefore for the efficient root-to-shoot translocation of B (Takano et al., 2002, Takano et al., 2010). Under limited B conditions, AtBOR2 is crucial for the transfer of B to cell walls in roots, allowing continued root elongation (Miwa et al., 2013). AtBOR4 effluxes B out of roots under toxic B conditions (Miwa et al., 2014). These pioneering discoveries in *A. thaliana* have paved the way for further studies advancing our understanding of B transport in other plant species (Chatterjee et al., 2014, 2017; Durbak et al., 2014; Hanaoka et al., 2014; Leonard et al., 2014; Zhang et al., 2017; Shao et al., 2018).

In contrast to the state-of-the-art progress made on B transport mechanisms, our current knowledge on the spectrum of molecular and physiological B-deficiency responses or the genetic and molecular basis of B-efficiency in *A. thaliana* is very scarce. Genetic analyses of physiological responses to low B stress in *A. thaliana* are restricted to a few studies considering only five different *A. thaliana* accessions (Zeng et al., 2007; Zeng et al., 2008; Huai et al., 2018). Analysis of B uptake, biomass production, seed yield and other B-dependent traits in four *A. thaliana* accessions defined “1938” and “Ler” as B-efficient and “1961” and “Col-4” as B-inefficient (Zeng et al., 2007). A recombinant inbred line population derived from a cross between “Ler” and “Col-4” subsequently resulted in the mapping of five B efficiency QTL (Zeng et al., 2008). More recently, At1g78860, encoding a curculin-like (mannose-binding) lectin family protein, was suggested to be the underlying gene for the B-efficiency QTL AtBE1-2 (Huai et al., 2018). How this protein can functionally impact on B efficiency has not yet been resolved. Further genes and proteins determining spatiotemporally variable B efficiency traits in *A. thaliana* have yet to be identified. As successful approaches in other species such as *Brassica napus* have demonstrated (Zhang et al., 2014; Pommerrenig et al., 2019), a valuable prerequisite for the discovery of B-efficiency QTL in *A. thaliana* would be a detailed overview of the natural variation in B-efficiency amongst diverse accessions and isolation of accessions with contrasting tolerance to B-deficiency. However, to date, detailed and systematic studies investigating natural variation in B-efficiency and B-deficiency tolerance in *A. thaliana* are lacking.

In this study, we comprehensively screened a genetically diverse collection of 185 *A. thaliana* accessions from across the world with the aim to identify and characterise natural variation in physiological, ionomic and molecular traits related to B-deficiency tolerance. Different manual and automated phenotyping technologies and cultivation systems allowed us to unravel and characterise 1) B-deficiency sensitive root and shoot growth traits, 2) morphological and molecular biomarkers for the detection of early B-deficiency, 3) highly B-deficiency tolerant and sensitive accessions, and 4) physiological traits that contribute to B-efficiency. The comparison of root and shoot system architecture and growth traits, elemental analyses and B transporter transcript abundance between accessions grown under different exogenous B conditions, combined with analysis of phylogenetic relationships, suggested that independent B efficiency strategies have evolved in individual and groups of *A. thaliana* accessions.

## Materials and Methods

### Plant material and growth

A total of 185 *A. thaliana* accessions were selected based on their expected genotypic variation and on their diverse origin representing sites expected to vary greatly in soil B availability. *Atnip5;1* knock-down mutant (SALK_122287; NASC: N622287) seeds were obtained from The European Arabidopsis Stock Centre (Loughborough, United Kingdom) and used as a control for B-deficiency treatments.

### Plant cultivation on soil-substrate in an automated plant phenotyping system

The automated phenotyping experiment took place from 23/03/2016 to 13/04/2016 at the IPK Gatersleben (51° 49’ 28.9” N, 11° 16’ 44.9” E). This was performed using an automated plant transport and imaging system (IPK automated plant phenotyping system for small plants) situated in a controlled environment plant growth chamber (Junker et al., 2015). Growth chamber conditions were set to long-day conditions (16 h day/8 h night) at 20/18 °C, 60/75% relative humidity, and ∼180 μmol light intensity light intensity. A total of 384 plant carriers were used, each consisting of 12 growth units of 4x4 cm (length x width) arranged in a 3x4 pattern (Fig. **1**). Carriers were arranged in 48 blocks of eight carriers situated across 12 conveyor belt lanes (six on each side of the plant growth chamber). The 185 selected *A. thaliana* accessions were represented across twelve biological replicates per accession and treatment (B-deficient; “-B” and B-sufficient; “+B”). Accessions were each distributed across six carriers, three of which were subjected to -B (< 0.1 mg-B kg-substrate^-1^) conditions, and three of which were subjected to +B conditions (∼2.4 mg B / kg substrate). On each carrier of 12 growth units, four plants of each of three accessions were cultivated. Replicate carriers were distributed throughout the system using a randomised block design.

**Fig. 1.**
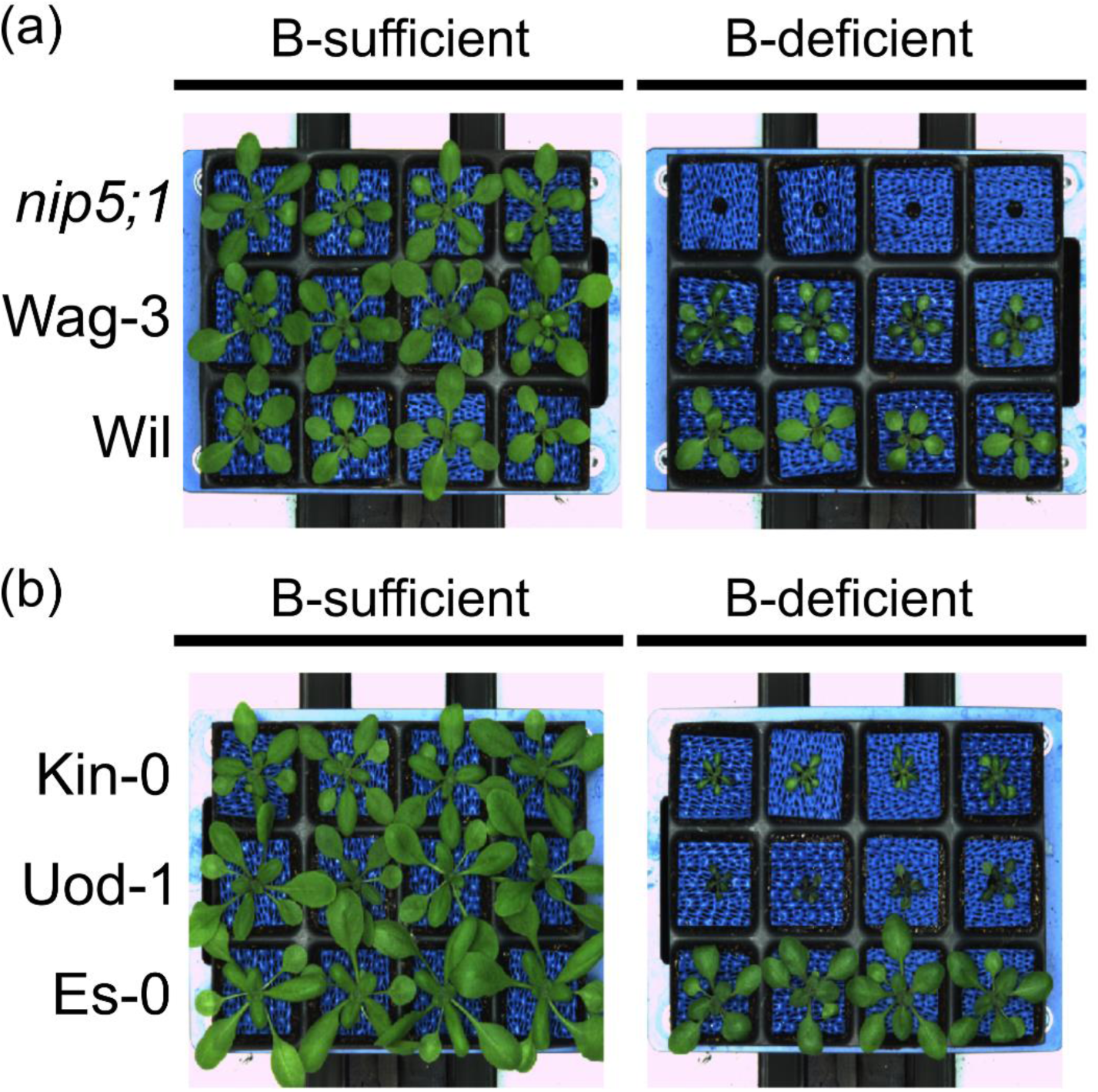
Example carriers used in automated phenotyping experiment. Images taken 20 days after sowing. Each of the 12 growth units per tray are 4x4 cm. Blue mats under rosettes used to increase contrast for automatic imaging and phenotype. The B transporter mutant *nip5;1* was included in trays in (a) as a control for the B-deficiency treatment: in such conditions, no growth was observed after germination. Accessions in (b) represent the range of genotypic performance observed, where Es-0 is a B-efficient accession, Uod-1 is a B-inefficient accession, and Kin-0 is a highly B-inefficient accession. Images were taken 20 DAS.

Growth substrate used was a B-free (< 0.1 mg-B kg-substrate^-1^) white-peat volcanic clay mixture (from here named zerosoil-substrate) with a dry matter of ca. 30%. Zerosoil-substrate was supplemented with 0.5% CaCO_3_ and 0.3% CaO (w/w) by mixing 20 kg batches of soil with 1 L of CaCO_3_ (100 g L^-1^) and 1 L of CaO (60 g L^-1^) in a clean cement mixer (Lescha Typ SBM P 150L) for 10 min. Thereafter, zerosoil-substrate was manually mixed every second day to allow even drying of the substrate. Around 1-2 weeks after preparation, clumps were sieved out. The following 100x micro- and 100x macronutrient stock solutions were prepared: NH_4_NO_3_ 60 g L^-1^, KH_2_PO_4_ 40 g L^-1^, K_2_SO_4_ 6 g L^-1^, MgSO_4_ 20 g L^-1^, CuSO_4_ 1.3 g L^-1^, ZnSO_4_ 1.3 g L^-1^, MnCl_2_ 4 g L^-1^, (NH_4_)_6_Mo_7_O_24_ x 4 H_2_O 8.5 mg L^-1^, NaFeEDTA 0.7 g L^-1^ and H_3_BO_3_ 1.4 g L^-1^. From these stocks, an 8x-concentrated nutrient solution was prepared for all nutrients excluding B and 250 mL were mixed per 1 kg sieved zerosoil-substrate in a cement mixer with a pressure sprayer. No H_3_BO_3_ stock was applied to zerosoil-substrate used for the B-deficient treatment. For the B-sufficient treatment, the H_3_BO_3_ stock solution was used to reach a final soil concentration of 2.5 mg B kg-soil^-1^. Each 4x4 cm growth unit was filled with ca. 17 g processed zerosoil-substrate.

Seeds of the 185 *A. thaliana* accessions were spread out on wet filter paper and left to pre-germinate in the dark at 20°C overnight, and then were sown out onto the zerosoil-substrate in each growth unit. In order to initiate germination, carriers underwent a stratification period of 3 days at 5°C in the dark (75% relative humidity) following seed sowing. After this, carriers were transferred to the plant growth chamber. Conditions were set to long-days (16h day/8h night) at 16/14°C, 75% relative humidity, for two days at 120 μmol light intensity and for two days at 180 μmol light intensity. After this, the temperature was increased to 20/18°C, with all other conditions kept the same. Plants were watered every day using an automated weighing and watering system to readjust to ca. 70% field capacity using ultrapure Milli-Q water (Merck, Milli-Q IQ7000) pre-treated with the B-chelator Amberlite IRA-743 (3 g L^-1^; Sigma) to prevent any B contamination.

Between 5 and 16 days after sowing (DAS), but excluding 11 DAS, top view images of all individual plants were taken daily in the visible range of the light spectrum (VIS), and of static fluorescence signals (FLUOR) as described previously in Junker et al. (2015). At the end of the experiment (20 DAS), shoot fresh weight (FW) of three pools of three representative plants per accession was measured by cutting the shoot directly above the substrate level and weighing. Dry weights were additionally quantified for each pool after drying the material at 60°C for 5 days. The relative shoot B-use efficiency indices (BEI_shoot_) of the 185 *A. thaliana* accessions were calculated as per equation [1]:

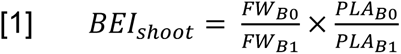

where FW_B0_ is the mean plant pool FW under -B conditions, FW_B1_ is the mean plant pool FW under +B conditions, PLA_B0_ is the mean projected leaf area in -B conditions based on fluorescence imaging (pixels^2^), and PLA_B1_ is the mean projected leaf area in +B conditions.

Relative growth rates (RGR) were calculated on both a daily basis and in three day intervals between 9 and 12 DAS, and 13 and 16 DAS, as per equation [2]:

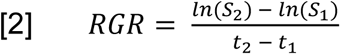

where ln is the natural logarithm, S_1_ is the size at time one, S_2_ is the size at time two, t_1_ is time point one and t_2_ is time point 2.

### Image and data processing

The Integrated Analysis Platform (IAP) was used for processing and automated feature extraction of images taken by the automated plant phenotyping system (Klukas et al., 2014). The multi-sensor setups at IPK (VIS, FLUO, NIR, FluorCam) supported the assessment of phenes corresponding to plant architecture, plant colouration, and levels of fluorophores, as well as efficiency of photosystem II. A total of 54 automatic imaging phenes were captured in the main database. Automated phenotyping data logged before day 9 were removed due to large amounts of missing data (plants were too small to be detected by the imaging system at earlier stages), and after day 16 because of overlapping rosettes. Across all 4,608 initially experimental units, 258 were removed completely from further analysis due to plants having not germinated or having died during the experiment, or due to the existence of more than one plant per growth unit. Data were removed from an additional 16 experimental units on specific measurement days due to poor imaging data on that day, for example due to plants not being visible. Individual trait outliers were determined separately for -B and +B datasets, and were defined as values lying above five standard deviations above the mean or below five standard deviations below the mean, for each trait measured on each day. In total, 1,120 values were removed across -B data, and 1,224 values were removed across +B data. In total, 1,629,632 values were retained across genotypes, treatments, measurement days and traits from the automated phenotyping experiment.

### Root phenotyping of *in vitro* cultivated plants

Square Petri dishes (120 x 120 x 17mm, vented; Greiner) were used as growth containers for the *A. thaliana in vitro* cultures (Fig. **2**). The experiment included the 185 selected accessions represented across five plants per Petri dish per accession and treatment (-B and +B). Seeds of each accession were surface sterilised and then cold stratified at 4°C for two days. They were then subjected to a pre-culture for five days in an autoclaved, modified half-strength Murashige and Skoog (½ MS) solidified medium comprising 625 µM KH_2_PO_4_, 750 µM MgSO_4_, 1.5 mM CaCl_2_, 9.4 mM KNO_3_, 1 mM NH_4_NO_3_, 75 µM FeNa_2_EDTA, 0.055 µM CoCl_2_, 0.05 µM CuSO_4_, 50 µM MnSO_4_, 0.5 µM Na_2_MoO_4_, 15 µM ZnSO_4_, 2.5 µM KI, 1 mM MES, 0.5% Sucrose, pH 5.5 (KOH), 1% Phytagel (Sigma). No B was specifically added to pre-culture media but nor was it specifically excluded. After cold stratification/pre-culture, five seedlings per accession were carefully transferred with forceps to autoclaved, solidified main-culture media with the same composition as the pre-culture medium but additionally containing 0.2 µM boric acid (-B) or 100 µM boric acid (+B). All main-culture media used was treated overnight with the B-chelator Amberlite IRA-743 (3 g/L) prior to autoclaving to remove all traces of B from the media. All Amberlite was sieved out of the growth media before autoclaving. The above-described boric acid concentrations were supplied to the media after Amberlite treatment and autoclaving, using separately autoclaved boric acid stock solutions.

**Fig. 2.**
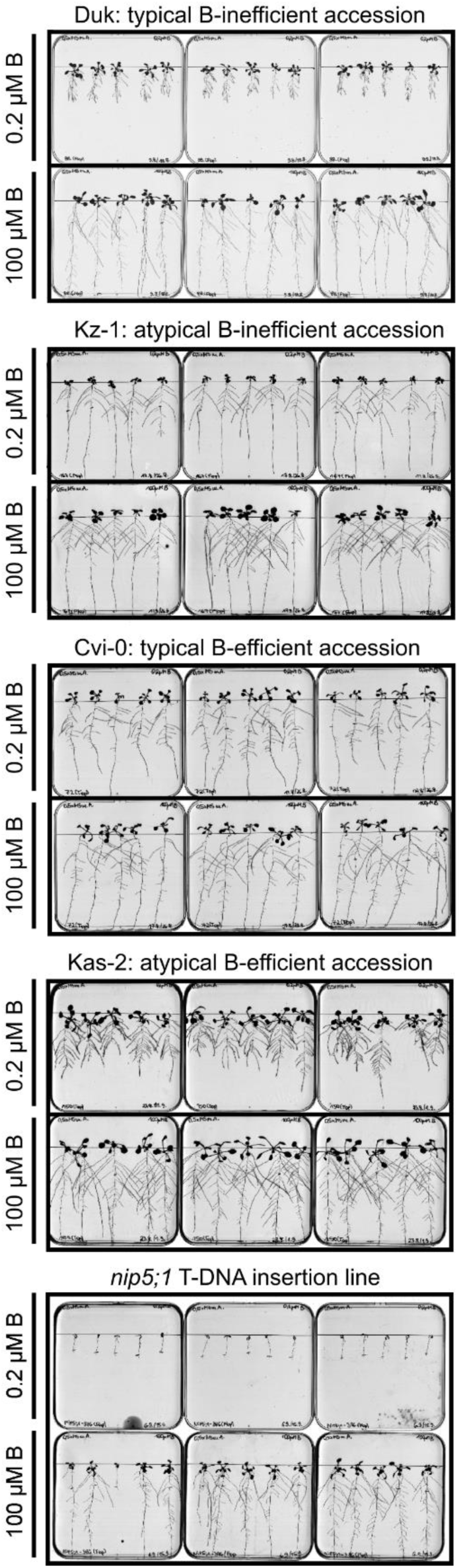
Example Petri dishes (12x12 cm) as used in *in vitro* root phenotyping experiments. Petri dishes contained either sufficient (100 µM) or deficient (0.2 µM) B concentrations. Accessions determined as B-efficient or B-inefficient in terms of shoot growth are shown; in each case, it is indicated whether root performance under B-deficient conditions is typical or atypical of the respective accession grouping. The B transporter mutant *nip5;1* was included in Petri dish experiments as a control for the B-deficiency treatment.

Petri dishes were wrapped around the vented edges with Micropore tape and placed vertically in a Percival Scientific CU chamber (CLF Climatics) with a photoperiod of 16 h light at 22°C (120 μmol m^−2^ sec^−1^) and 8 h dark at 19°C. All plates were re-positioned every second day to avoid positioning effects in the growth chamber. After 10 days of growth on the main-culture media, plates were scanned using an Expression 11000XL scanner (Epson). Shoots were then carefully separated from roots with forceps and biomass (fresh weight) of pools of 3-5 plants per accession and treatment was determined. After 3 days drying at 65°C, the dry weights of the same pools were determined. Root systems were combed to prevent any overlap between roots prior to scanning. Primary root length (PRL), total lateral root length (TLRL), average lateral root length (ALRL) and lateral root number (LRN) were determined by analyzing the scanned images using EZ-Rhizo software (Shahzad et al., 2018). The relative root B-use efficiency indices (BEI_root_) of the 185 *A. thaliana* accessions were calculated as per equation [2]:

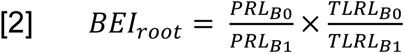

where PRL_B0_ is the mean primary root length under -B conditions, PRL_B1_ is the mean primary root length under +B conditions, TLRL_B0_ is the mean total lateral root length in -B conditions, and TLRL_B1_ is the mean total lateral root length in +B conditions. Twelve plants per accession and condition were used for the calculation of the mean root trait values.

### Mineral elemental analysis

Shoot ionomes of plants grown on both B treatments in zerosoil-substrate in the automated plant phenotyping system were determined for three pools of three representative biological replicates of 43 selected *A. thaliana* accessions after harvest at 20 DAS. Accessions were selected based on their BEI_shoot_ scoring (equation [1]): 17 B-efficient accessions of the 90^th^ percentile (Var2-1, Hn-0, Yo-0, Cha-0, Tha-1, Db-0, Cal-0, Ty-0, Sorbo, Cvi-0, Ca-0, CUR-3, TOU-A1-116, Es-0, Zü-1, N7, Eden-1), 5 B-intermediate efficient accessions with BEI_shoot_ above 0.4 (Kn-0, Kas-2, Liarum, S96, In-0), three B-inefficient accessions (Col-0, Limeport, Uod-1) and 18 highly B-inefficient accessions of the 10^th^ percentile (Bay-0, Duk, Gel-1, Hau-0, Hovdala-2, Kin-0, Kz-1, Nz1, Pa-1, Ri-0, Rld-2, Rmx-A02, Tsu-1, Tu-0, Ull2-3, Van-0, WAR, Ws). Plant material was sampled and dried at 65°C for one week. Approximately 5-10 mg of plant dry matter was digested in nitric acid (HNO_3_) using a high-performance microwave reactor (UltraClave IV; MLS GmbH, Leutkirch, Germany). Elemental analysis was performed with a sector field high-resolution ICP-MS (Element 2, Thermo Fisher Scientific, Waltham, MA, USA) using software v.3.1.2.242 (Eggert and von Wirén, 2013). The elements B, Na, Mg, P, S, K, Ca, Mn, Fe, Cu and Zn were quantified.

### RNA extraction, cDNA synthesis, and real-time quantitative PCR

Approximately 10-40 mg *A. thaliana* plant material from the accessions as selected above was harvested from the plant roots, immediately frozen in liquid nitrogen and ground with a pestle and mortar in liquid nitrogen. Total RNA was extracted using NucleoSpin RNA Plant Kits and DNase I treatment according to the manufacturer’s instructions (Macherey-Nagel, Germany). Complementary DNA (cDNA) was synthesised from 1000 ng total RNA per sample using MuLV-Reverse Transcriptase (Fermentas, Germany) in a total volume of 20 µl and diluted to 1:20 with nuclease free water. RT-qPCR was performed in a 384 well thermocycler (CFX384 TouchTM Real-Time PCR Detection System, Bio-Rad) using the GoTaq qPCR Mastermix (Promega, USA). Per reaction, 2 µl diluted cDNA were used. Three identically treated biological replicates were analysed. The thermocycler protocol was as follows: 3 min at 95°C, followed by 45 cycles of 10 s at 95°C, and 50 s at 58 or 60°C. For the generation of a standard curve, aliquots of the above-described cDNA samples of diverse tissue pools of the corresponding growth conditions were used in a mixture. These mixtures were serially diluted (1 to 1, 1 to 2, 1 to 4, 1 to 8, 1 to 16, 1 to 32, and 1 to 64) with nuclease free water to generate standard curve templates and to determine PCR efficiencies for each primer pair. Assays producing PCR efficiencies > 80% < 115% were used for expression data analyses. Relative expression of At*NIP5;1*, *AtNIP6;*1, and *AtBOR1* using the reference gene *AtEF1a* was determined by qPCR as previously published (Takano et al., 2006; Tanaka et al., 2008) using primers summarised in Table **S1**.

### Phylogenetic and *in silico* parameter analysis

A maximum parsimony phylogenetic tree of the 185 accessions was generated based on 214,051 Single Nucleotide Polymorphisms (SNPs; Horton et al., 2012, Atwell et al., 2010) using the R package “phangorn” (Schliep, 2011) and implementing the parsimony ratchet (Nixon, 1999). Accessions were manually assigned to one of 11 clades within the tree. B-efficient genotypes were plotted onto a geographic map using the function geom_map in the package “ggplot2”, using the map “world_coordinates” as a background. Accessions were colour-coded to the phylogenetic clade determined in the previously described analysis. *k*-means clustering was used to group accessions by shared behavior across multiple shoot and root traits using the stats package in the software R. *k*-means clustering was based on a total of 38 traits, which were all ratios of the mean accession performance under -B compared to +B conditions. Of the 54 phenes retained from the automated phenotyping system, 11 were selected to represent shoot development for *k*-means clustering. These comprised projected leaf area, plant border length, branchpoint count, hsv h brown:green pixel ratio, hsv h yellow:green pixel ratio, hull area, hull length, hull points count, leaf count, mean leaf length, and mean leaf width. The remaining 43 phenes were excluded due to being aliased with another included phene (18), having limited variation either between genotypes or between growth conditions (15), concerning a non-plant relevant shape (e.g. rectangle area; 6), or contradicting with other measured phenes (4). Each phene was included at days 9, 12 and 16. The phene “hsv h brown:green pixel ratio” was not included at days 9 and 12, and “branchpoint count” was excluded at day 9, each due to missing data for most genotypes. Relative growth rates in the period from day 9-12 and day 13-16, as well as dry weight at experiment end, were additionally included, as were the four measured root traits and DW of shoots from the *in vitro* experiment. The optimal number of *k*-clusters to use was determined using the fviz_nbclust function within the package “factorextra”, considering both the “Elbow Method” and “Silhouette method”. Twelve *k*-clusters were selected as the best representation of the data. Following assignment of accessions to a *k*-cluster, trait means for each *k*-cluster were recalculated, centred and scaled for generation of Fig **9**.

### Proteoform Analysis

Amino acid sequence variations in the B transporters AtNIP5;1 (At4g10380), AtNIP6;1 (At1g80760), AtNIP7;1 (At3g06100), and AtBOR1(At2g47160) of 1135 *A. thaliana* accessions were analyzed using the SNP★ar tool (http://141.48.9.71). In the SNP★ar tool, the transcript identifier of the B transporters was selected and searched for the SNPs in the database available for the *A. thaliana* accessions. Changes in the nucleotide sequences that result in non-synonymous amino acid substitutions in the four above-mentioned B transporters were identified and extracted and their frequency was analysed amongst all 1135 accessions of the database and the available accessions present in this study.

## Results

### Highly B-efficient *Arabidopsis thaliana* accessions identified by analysing natural variation in shoot morphology under B-deficient growth conditions

A population of 185 *Arabidopsis thaliana* accessions were grown on zerosoil-substrate under B-sufficient (2.5 mg B /kg) or B-deficient conditions (< 0.1 mg B /kg) in an automated phenotyping facility. Plant growth and development were documented and quantified on a daily basis by top view imaging using a fluorescence and a visible light camera from 9 to 16 days after sowing (DAS). The cultivation was optimised for a camera-based quantification of the projected leaf area, including through laying blue mats under plant rosettes for higher contrast imaging (Fig. **1**). Within the experimental duration, most accessions showed clear B-deficiency symptoms compared with plants in the control treatment (Fig. **1a,b**). This was primarily visible as reduced shoot growth, leaf curling, and darker green leaf color. Whilst most accessions continued to grow in size with time, plants subjected to B-deficiency grew at a much slower pace. This is reflected by the projected leaf area (PLA; Fig. **3a**,**b**), which correlated strongly with shoot dry weight at experiment end (20 DAS, Fig. **3c**, Fig. **S1**), thereby validating the experimental setup and analysis pipeline. Average PLA differed clearly between deficient and sufficient B conditions from around 12 DAS (Fig. **3b**). The growing difference in PLA between B-sufficient and B-deficient plants demonstrated that the longer plants were exposed to B-deficiency, the more the growth inhibition manifested.

**Fig. 3.**
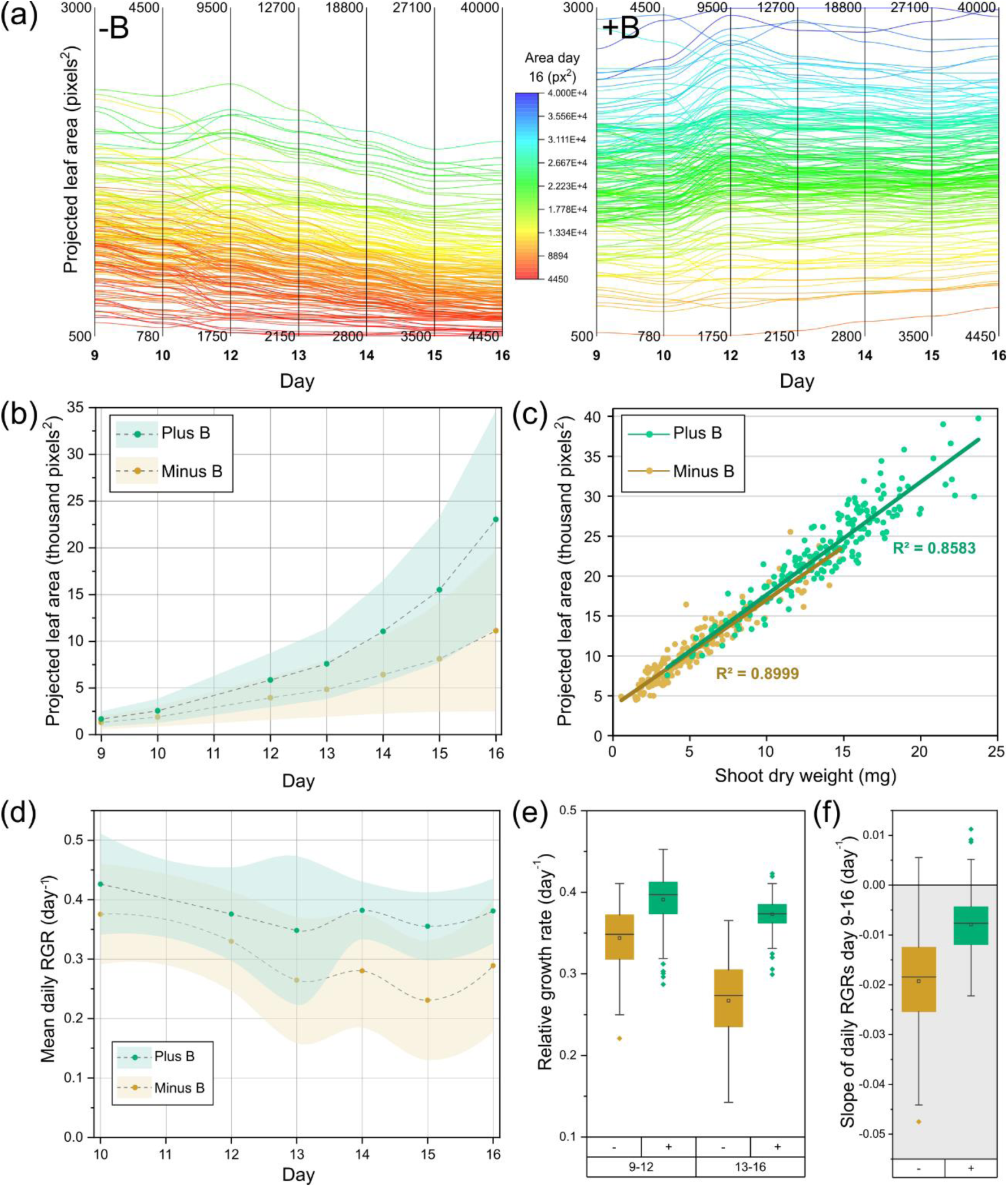
Growth of accessions in the automated phenotyping experiment over time. (a) Parallel plots showing the change in projected leaf area (PLA) over time in deficient (-) or sufficient (+) B growth conditions. Each line represents an accession. (b) Mean average growth performance of accessions over time. Dashed lines and dots represent mean average PLA across genotypes on each day in each B treatment. Shaded areas surrounding lines represent two standard deviations on each side of mean values. PLA between measurement days inferred with Akima Spline smoothing. (c) Correlation between shoot dry weight at day 20 (experiment end) and PLA at day 16 (last imaging day) for both B-sufficient (Plus B) and B-deficient growth conditions. (d) Change in daily relative growth rate (RGR) of accessions in B-sufficient (Plus B) or B-deficient (Minus B) conditions. Dashed lines and dots represent mean average daily RGR across genotypes on each day in each B treatment. Shaded areas surrounding lines represent two standard deviations on each side of mean values. RGRs between measurement days inferred with Akima Spline smoothing. (e) RGRs in three day intervals from 9-12 and 13-16 DAS across genotypes between B-sufficient (+) and B-deficient (-) conditions. (f) The slope of daily RGRs across genotypes between B-sufficient (+) and B-deficient (-) conditions, where a more negative slope represents a faster reduction in RGR with time. For box plots, boxes represent the 25^th^ to 75^th^ percentiles and whiskers represent the range within 1.5 times the interquartile range (IQR) with any outliers shown.

On average, accessions suffered a 50% reduction in PLA in B-deficient compared to B-sufficient conditions by day 16. Thirteen of the 185 accessions maintained above 80% of their mean B-sufficient PLA when grown in B-deficient conditions. Interestingly, the accession Var2-1 was ca. 28% bigger in B-deficient than B-sufficient conditions at day 16, and was the only accession to be bigger when exposed to B-deficiency. However, Var2-1 was in only the 10^th^ percentile for PLA in B-sufficient conditions. It is therefore a small plant in general which probably has a low nutrient demand, and as such may not be the most B-efficient accession in this dataset. Of the accessions in the 90^th^ percentile for PLA in B-sufficient conditions, none maintained as much as 80% of their mean B-sufficient PLA when grown in B-deficient conditions. However, the 90^th^ PLA percentile accession, Es-1, was able to maintain 77% performance under B-deficiency, and thus may be considered B-efficient. Accession relative growth rates (RGRs) were calculated on both a daily (Fig. **3d**) and a 3-day (Fig. **3e**) basis. Daily RGRs generally decreased between 9 and 16 DAS across genotypes grown in both B conditions, as reflected by the slope of daily RGRs by day (Fig. **3f**). However, the slope was more negative for plants grown in B-deficient conditions. As such, a clearly lower daily RGR was apparent across accessions from 14 DAS (Fig. **3d**). Similar effects were observed for the 3-day RGRs, with a bigger difference in RGR between B-treatments observed in the second half of the imaging period (13-16 DAS; Fig. **3e**).

Accessions were assigned a shoot B efficiency index (BEI_shoot_) based on equation [1], which considers both plant FW at experiment end (20 DAS) and PLA at the final imaging day (16 DAS). The average BEI_shoot_ was 0.230 and ranged from the minimum value of 0.016 (Kz-1) to the maximum value of 1.521 (Var2-1). Accessions were grouped into four growth groups. Group 1 comprises 19 B-efficient accessions in the 90^th^ BEI_shoot_ percentile with BEI_shoot_ values all higher than 0.479 (Var2-1, Hn-0, Yo-0, Cha-0, Tha-1, Db-0, Cal-0, Ty-0, Sorbo, Cvi-0, Ca-0, CUR-3, TOU-A1-116, Es-0, Zü-1, N7, Eden-1, Ge-1, Com-1). Group 2 comprises 11 B-intermediate efficient accessions with BEI_shoot_ values above 0.400 (Kn-0, Bl-1, Kas-2, Bs-1, Liarum, S96, In-0, Kondara, Je-54, Zdr-1, Co). None of the B-efficient or B-intermediate efficient accessions suffered more than a 40% reduction in PLA under B-deficient compared to B-sufficient growth conditions. The six genotypes Var2-1, Yo-0, Cha-0, Tha-1, Hn-0 and Db-0 suffered no more than a 20% reduction in either PLA, FW, or DW under B-deficient compared to B-sufficient growth conditions, and are therefore suggested to be highly B-deficiency tolerant accessions with respect to shoot growth. Of these six accessions, three also had a higher shoot DW in B-deficient compared to B-sufficient growth conditions: the shoot DWs of Yo-0 and Hn-0 were 3 and 2% heavier under B-deficiency, and the shoot DW of Var2-1 was 33% heavier, suggesting even a preferential growth in low B environments for this latter accession. Amongst the remaining accessions, we defined the 136 accessions with BEI_shoot_ over the 10^th^ percentile as B-inefficient lines (Group 3), and the 19 accessions with BEI_shoot_ less than the 10^th^ percentile (BEI_shoot_ < 0.043) as highly B-deficiency sensitive accessions (Group 4; Hovdala-2, Pa-1, Rmx-A02, Ws, Hau-0, Tu-0, Tsu-1, Ri-0, Alc-0, Kin-0, Nz1, Ull2-3, Van-0, Bay-0, WAR, Duk, RLD-2, Gel-1, Kz-1). Representative B-efficient (Es-0) and B-inefficient (Wag-3, Wil, Kin-0 and Uod-1) accessions as well as the *atnip5;1* knockout mutant are displayed from both B treatments in Fig **1** (**a,b**). *Atnip5;1*, which did not develop beyond germination under B-deficient conditions, served as a biological control genotype to ensure establishment of B-deficient growth conditions.

### Shoot architecture and colour-related phenes serve as early predictors for the classification of B-sufficient and B-deficient growth conditions

Of the 378 traits measured over the seven imaging days of the automated phenotyping experiment (54 phenes measured per day), 105 were more than 20% higher or lower in B-deficient compared to B-sufficient growth conditions on average across accessions. The majority of these (79) were apparent only from 13 DAS, with only 26 traits differing by more than 20% between B-deficient and B-sufficient growth conditions between 9 and 12 DAS. The area of the plant convex hull, defined as the minimum convex set that contains the plant rosette, generally followed PLA in terms of variation in the response to B-deficiency between accessions (Fig. **4**). However, whilst the number of hull points also increased with experimental duration for both treatments, differences between B conditions were clearly apparent from 9 DAS. This is earlier than the point at which differences in PLA or hull area became apparent, and so this trait could be used as an early marker for B-deficiency in plants. Similarly, B-deficient plants exhibited generally more branching than B-sufficient plants at 9 and 10 DAS, although this response was less apparent as the experiment continued (Fig. **4**). Among the imaging dataset were various colour-related phenes. Most notably, values of the RGB, HSV and LAB colour models were quantified, as were ratios of yellow to green and brown to green pixels. Colour-related traits were more represented at the first half of the imaging period, making up just over 23% of the traits differing by more than 20% between B treatments in this period compared to only 15% in the latter half of the imaging period. RGB green values were generally lower across all imaging days in B-deficient growth conditions, ranging from a 7% decrease at day 9 to an 11% decrease at day 16 (Fig. **4**). RGB green also correlated with BEI_shoot_ across all imaging days, with Pearson’s correlation coefficients ranging from 0.32 at day 9 to 0.58 at day 16 (all *p* < 0.001). Such a phene could be a useful measure for early detection of plant B-deficiency, assuming a detection system capable of differentiating B-deficiency from other deficiencies such as nitrogen could be engineered (Zhang et al., 2024). Mean L values of the LAB colour model and V values of the HSV colour model also followed similar patterns to RGB green, and so these phenes could also be fed into any model designed for early B-deficiency detection. Both the ratios of yellow to green and brown to green pixels also responded to B-deficiency. However, these responses appeared first later, typically from 15-16 DAS (Fig. **4**). Photosynthetic operating efficiency of all plants was measured 16 DAS using a PAM fluorometer with 590 µmol m^-2^ s^-1^ photosynthetically active radiation. Across all accessions, photosynthetic operating efficiency was on average ca. 5% higher in plants subjected to B-deficient compared to B-sufficient conditions (Fig. **S2**). However, no correlations were observed between this and BEI_shoot_ or BEI_root_.

**Fig. 4.**
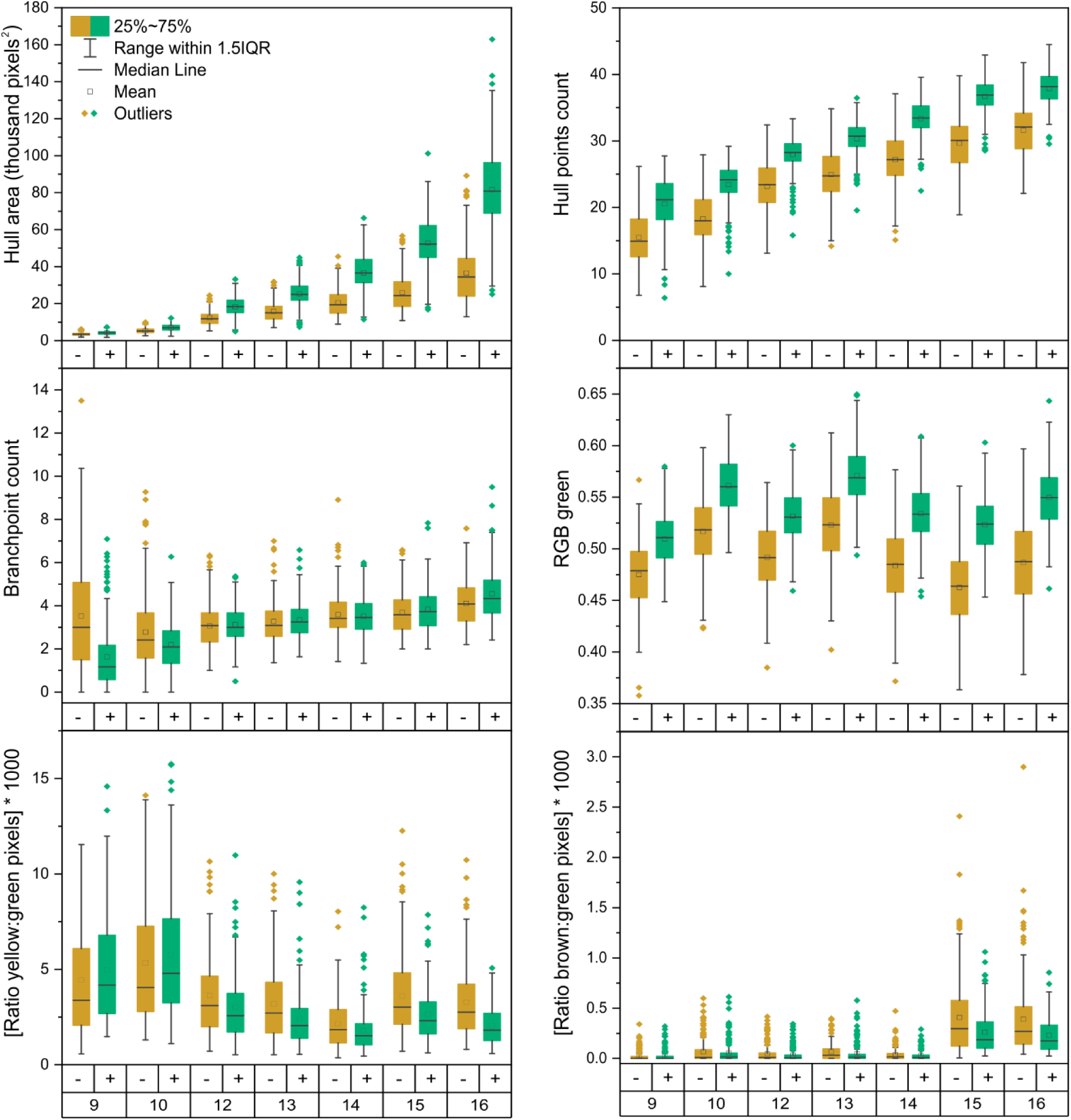
Change in three selected geometry and three selected colour phenes with time across accessions grown in either B-sufficient (+) or B-deficient (-) conditions. For all box plots, boxes represent the 25^th^ to 75^th^ percentiles and whiskers represent the range within 1.5 times the interquartile range (IQR) with any outliers shown.

### Boron deficiency reduced shoot B concentration but to a similar degree in both B-efficient and B-inefficient accessions

The shoot ionome of 43 accessions was determined using tissue harvested 20 DAS from the automated phenotyping experiment. Accessions were selected based on their BEI_shoot_. Boron-efficient accessions were represented by 17 of the 19 Group 1 (B-efficient) accessions and five of the Group 2 (B-intermediate efficient) accessions. B-inefficient accessions were represented by 18 of the 19 Group 4 (highly B-deficiency sensitive) accessions and three B-inefficient accessions of Group 3.

Boron-efficient accessions had generally higher B concentrations than B-inefficient accessions in both B-sufficient (11% higher) and B-deficient (40% higher) growth conditions. However, both groups were similarly affected by B-deficiency, with an average 84% reduction in shoot B concentration in B-efficient accessions grown in B-deficient conditions, compared to an average 87% reduction in B-inefficient accessions (Fig **5a**). Under B-sufficient growth conditions, shoot B concentration in B-inefficient accessions varied between 51.1 mg kg^-1^ DW (Tsu-1) and 79.9 mg kg^-1^ DW (WAR), and in B-efficient accessions between 55.9 mg kg^-1^ DW (Var2-1) and 91.7 mg kg^-1^ DW (Hn-0; Fig. **5a**). Shoot B concentration in B-deficient growth conditions varied in B-inefficient accessions between 5.1 mg kg^-1^ DW (Hau-0) and 15.3 mg kg^-1^ DW (Tsu-1), and in B-efficient accessions between 8.1 mg kg^-1^ DW (Kn-0) and 16.8 mg kg^-1^ DW (S96). Shoot concentrations of other mineral nutrients were generally not affected by B-deficiency across the B-efficient accessions (Fig. **S3**, **S4**). However, B-inefficient accessions exhibited lower Mg, P, Ca, Mn, Fe and Zn in B-inefficient accessions in B-deficient conditions, probably as a result of a general decline in plant health in such conditions. Interestingly, shoot K concentrations were lower in many of the B-inefficient accessions compared to B-efficient accessions in both B-sufficient and B-deficient conditions (Fig. **S3**). Across all accessions, shoot concentrations of S and Na were on average 47 and 24% higher in plants grown in B-deficient compared to B-sufficient growth conditions. Since concentrations of many nutrients were not affected by, or decreased, in response to B-deficiency, the increased concentrations of S and Na are unlikely to be a result of smaller plant size in B-deficient conditions. This therefore suggests that the uptake or the root to shoot translocation of these elements is reduced under B-deficiency.

**Fig. 5.**
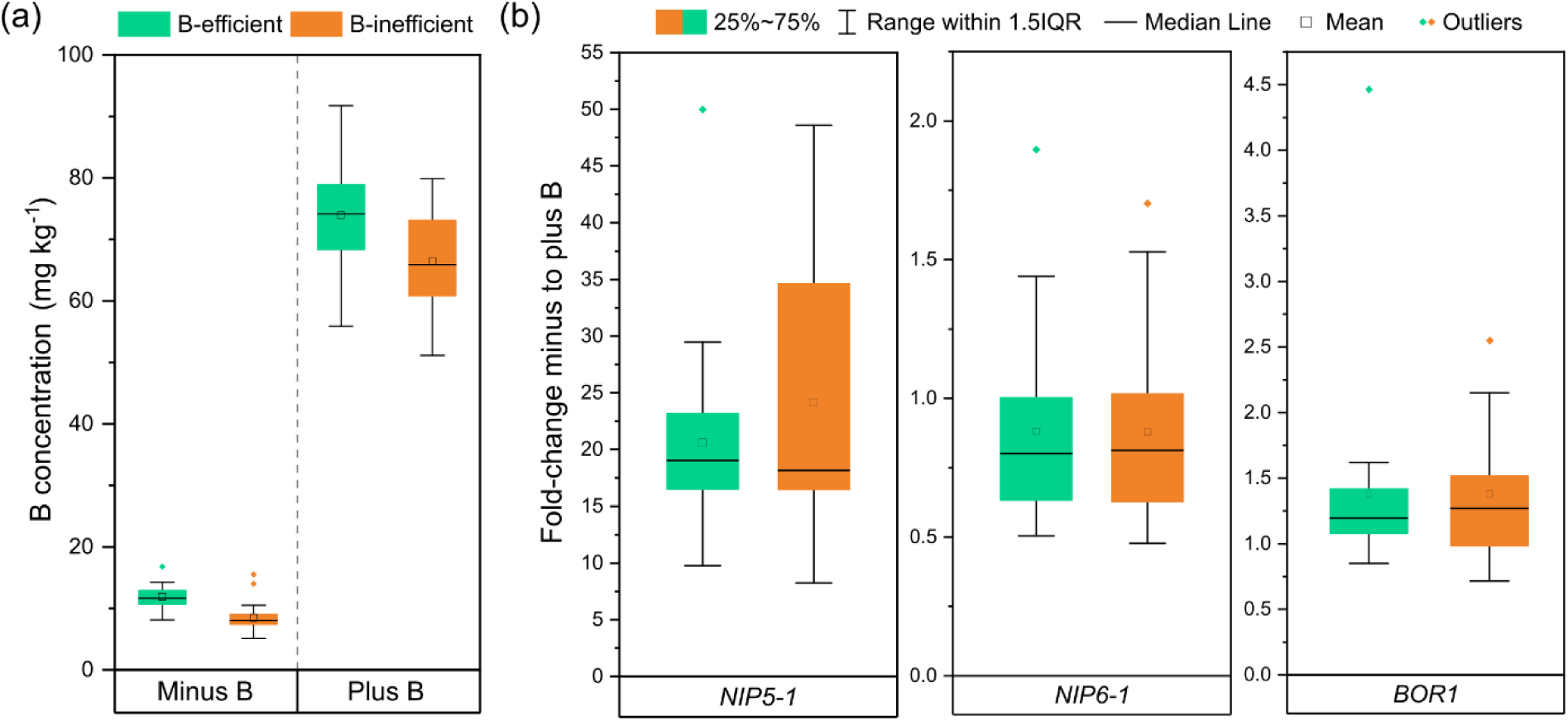
Shoot B concentration and B transporter expression profiles of 43 selected accessions. “B- efficient” boxes comprise data from 17 and 5 accessions defined as B-efficient and B-intermediate efficient, respectively, and “B-inefficient” boxes comprise data from 3 and 18 accessions defined as B- inefficient or highly B-inefficient, respectively (see Materials and Methods). (a) Shoot B concentration of three pools of three representative biological replicates of each accession of plants grown on both B treatments in zerosoil-substrate in the automated plant phenotyping system. (b) Fold-change in expression of *NIP5;1*, *NIP6;1* and *BOR1* in B-deficient (minus B) compared to B-sufficient (plus B) conditions of roots the same 43 accessions grown in the *in vitro* root phenotyping experiment. For all box plots, boxes represent the 25^th^ to 75^th^ percentiles and whiskers represent the range within 1.5 times the interquartile range (IQR) with any outliers shown.

### Shared and varied responses to B-deficiency observed between soil-substrate growth and *in vitro* growth systems

In addition to the soil-substrate cultivation experiment, we assessed the shoot growth response to B-deficiency of the 185 *A. thaliana* accessions in a Murashige-Skoog (MS)-supplemented agar-based Petri-dish cultivation system at an early developmental stage (15 DAS including 10 days exposed to contrasting B availabilities). The average shoot DW across accessions was 17% lower under B-deficient compared to B-sufficient conditions, clearly demonstrating a nutritional stress at this early seedling stage (Fig. **6a**). On average across accessions, the ratio of DW in B-deficient compared to B-sufficient conditions was 0.83. The average ratio of PLA in B-deficient compared to B-sufficient conditions as measured in the soil-substrate cultivation system after 10 days of growth in B-deficient conditions was comparable (0.76). The *in vitro* growth system can therefore be considered comparable to the soil-substrate system in its ability to induce B-deficiency in *A. thaliana*. Individual accessions were however not always identical in their ability to gain shoot biomass in these two contrasting growth conditions. The shoot DW of accession Var2-1 was 33% higher under B-deficient compared to B-sufficient growth conditions in the soil-substrate cultivation system, but 41% smaller in the *in vitro* growth system. Thirty-four percent of accessions in the top 20^th^ percentile of the DW ratio between B-deficient and B-sufficient growth conditions in the soil-substrate cultivation system were also in the top 20^th^ percentile in the *in vitro* system. Over 60% of accessions were in the top 50^th^ percentile for DW ratio in both systems, so a general shared genetic basis of B-efficiency in both systems was observed.

**Fig. 6.**
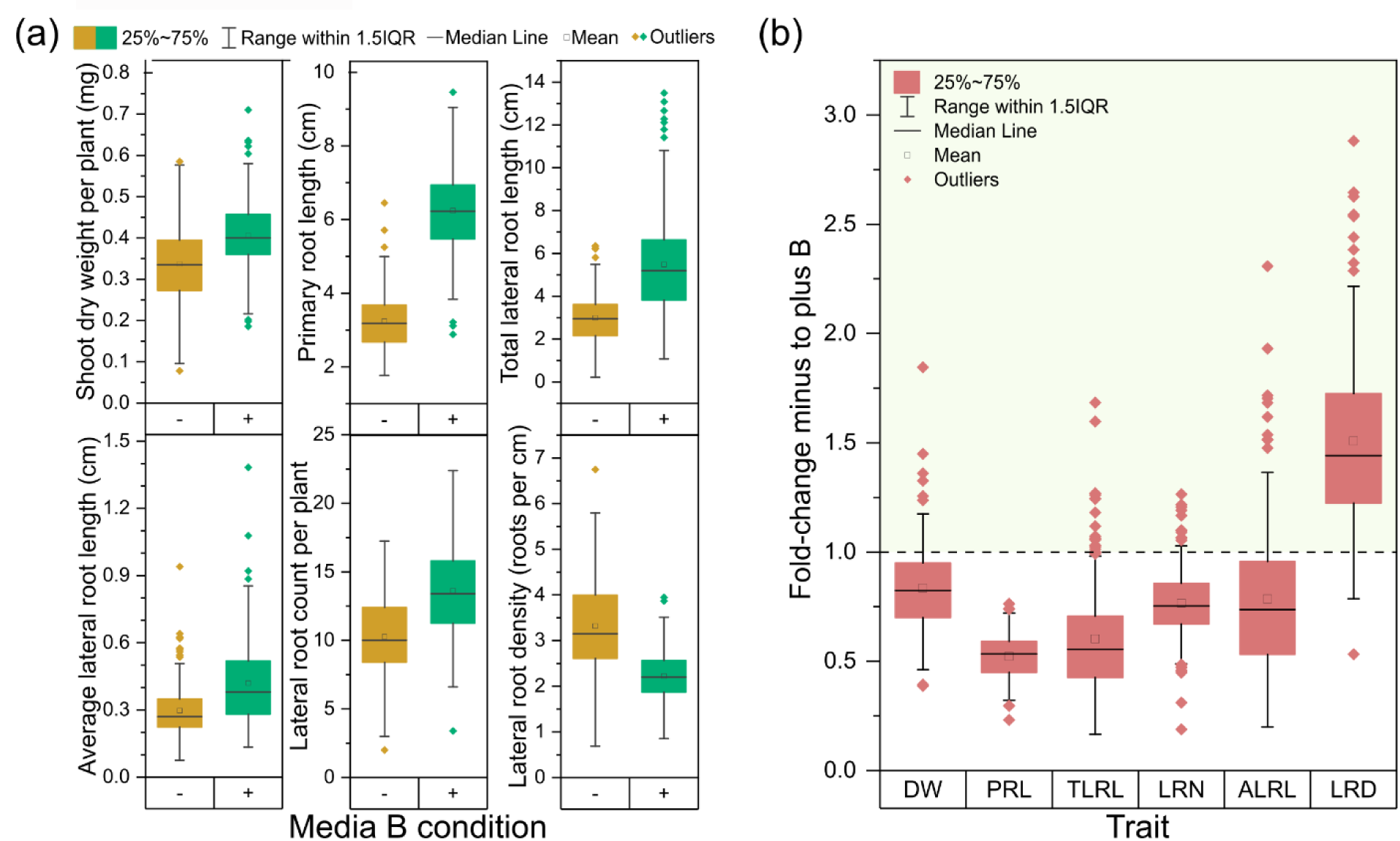
A summary of accession performance for traits measured in the *in vitro* root phenotyping experiment. (a) Accession variation in shoot dry weight and root system architecture phenes in B-sufficient (+) and B-deficient (-) conditions. (b) Fold-change for each phene for each accession of performance under B-deficient (-) compared to B-sufficient (+) conditions. Dashed line at 1 on the *y*-axis shown to indicate direction of accession response, where data above one indicate greater phene values under B-deficient (-) compared to B-sufficient (+) conditions. For all box plots, boxes represent the 25^th^ to 75^th^ percentiles and whiskers represent the range within 1.5 times the interquartile range (IQR) with any outliers shown.

### Primary root length was the root system architecture phene most strongly affected by B-deficiency, whilst increased lateral root length may be an adaptive response

We also assessed the root response to B-deficiency of the 185 *A. thaliana* accessions in the Petri-dish cultivation system. The length of the primary root (PRL) decreased in all accessions under B-deficiency, ranging from a 24% PRL reduction in N13 to a 77% reduction in Rak-2. The average PRL reduction across accessions was 48% (Fig. **6a,b**). Total lateral root length (TLRL) decreased on average by 40% under B-deficient growth conditions. However, TLRL of 16 accessions was higher in B-deficient than B-sufficient conditions, indicating a possible adaptive response. Two of these accessions, Eden-1 and Yo-0, were also in the 90^th^ percentile for BEI_shoot_. The minimum reduction in TLRL observed was 0.1% (Gy-0) and the maximum reduction was 83% (Tamm-2, demonstrating the high variability of this trait). An increase in lateral root number (LRN) was also observed in 15 accessions under B-deficiency, although only two of these were among the accessions with an increase in TLRL. In all other accessions, LRN decreased by an average of 26%. For average lateral root length (ALRL), 38 accessions exhibited an increase under B-deficiency, comprising 15 of the 16 accessions that exhibited an increased TLRL under B-deficiency, and seven of the accessions in the BEI_shoot_ 90^th^ percentile (Fig. **6b**). Other accessions exhibited on average a 35% reduction in ALRL, but over a 60% reduction was observed for 18 accessions in B-deficient conditions down to a maximum reduction of 80% for the accession, Hod. Finally, lateral root density (LRD), defined by the number of lateral roots per cm of primary root, showed generally higher values in B-deficient compared to B-sufficient conditions. This was due to the observed larger general decrease in PRL than LRN under B-deficiency across accessions. The average increase in LRD under B-deficiency was 51%, ranging up to more than a 100% increase for 18 accessions. Most of these 18 accessions were considered B-inefficient based on their BEI_shoot_ scores. However, one accession, Kas-2, was considered a B-intermediate efficient accession, with a BEI_shoot_ of 0.46.

Among the root traits which exhibited a decrease in B-deficient compared to B-sufficient growth conditions on average across accessions, PRL was most sensitive to B-deficiency followed by, in the following order: TLRL > LRN > ALRL. However, the ratio of TLRL under B-deficient compared to B-sufficient growth conditions was more highly correlated with shoot DW than PRL, with Pearson’s correlation coefficients of 0.47 and 0.35, respectively. This suggests that TLRL impacts more on shoot biomass formation than PRL under B-deficient conditions at an early growth stage, and adds further weight to the argument that TLRL may be an important adaptive trait contributing to B efficiency. Comparing results of the Petri-dish cultivation system with the soil-substrate system confirmed this interaction. The Pearson’s correlation coefficient between TLRL in agar and DW at 20 DAS in soil-substrate, both in B-deficient conditions, was 0.32, compared to only 0.18 between PRL and DW. The same comparisons in B-sufficient conditions yielded Pearson’s correlation coefficients of only 0.11 and 0.07, respectively, therefore confirming an important contribution of primary roots for growth in low B environments, but an even more important role of lateral roots.

Accessions were assigned a root B efficiency index (BEI_root_) based on equation [3], which considers the responses of both PRL and TLRL to B-deficiency. The average BEI_root_ was 0.32 and ranged from 0.054 (Rak-2) to 1.243 (DraIV 6-35). We classified accessions of the 90^th^ BEI_root_ percentile (> 0.52) as B-efficient in terms of growth in the Petri-dish cultivation system (19 accessions). Five accessions were in the 90^th^ percentile of both BEI_shoot_ and BEI_root_ and so were considered B-efficient in both growth environments. These were Cha-0, Eden-1, Tha-1, Hn-0 and Yo-0. Four accessions (Van-0, Kz-1, RLD-2 and Gel-1) were in the 10^th^ percentile of both BEI_shoot_ and BEI_root_ and so were considered highly B-deficiency sensitive in both growth environments. The Pearson’s correlation coefficient between BEI_shoot_ and BEI_root_ was 0.26. The phenes PRL, TLRL and ALRL under B-deficient conditions all correlated with BEI_root_ with Pearson’s correlation coefficients of 0.35, 0.38 and 0.33, respectively. Two accessions, Uod-1 and Hovdala-2, were in the 90^th^ percentile of BEI_root_ but in the 10^th^ percentile of BEI_shoot_. In contrast, none of the accessions in the 90^th^ percentile of BEI_shoot_ were in the 10^th^ percentile of BEI_root_. This indicates that whilst a large root system in B-limiting conditions does not necessarily correlate with good shoot performance, accessions with high shoot performance under B-deficiency benefitted from root system architecture traits that were not completely inhibited by B-limitation.

### Accessions with contrasting tolerance to B-deficiency show no differences in the expression of B transporter genes

Transcript abundance of the three B transporters *AtNIP5;1*, *AtNIP6;1* and *AtBOR1*, which were demonstrated to be important for B transport (Takano et al., 2002; Takano et al., 2006, Tanaka et al. 2008), was determined in roots of seedlings of the same 43 accessions selected for shoot ionome analysis (Fig. **5b**, Fig. **S5**). Over the tested accessions, the range of relative expression levels was on average relatively narrow for *AtNIP5;1* (3.8- and 3.5-fold variation under B-sufficient and B-deficient conditions, respectively) and *AtBOR1* (4.8- and 4.6-fold variation, respectively) but much wider for *AtNIP6;1* (30.6- and 23.6-fold variation, respectively). *AtNIP5;1* was strongly upregulated in all tested accessions under B-deficient compared to B-sufficient growth conditions. Upregulation varied from 8-fold to 50-fold between accessions, with an average of 22-fold upregulation. Despite this response to B-deficient growth conditions, the average level of upregulation did not significantly differ between the B-efficient and B-inefficient accessions (*p* = 0.24). The average upregulation of *AtNIP5;1* across the 22 analysed B-efficient accessions was 20.6-fold and across the 21 analysed B-inefficient accessions was 24.2-fold. Both groups contained accessions with relatively higher (Kas-2, B-efficient, 50.0-fold; Duk, B-inefficient, 48.6-fold) and lower (Ca-0, B-efficient, 9.8-fold; Rld-2, B-inefficient, 8.3-fold) upregulation. *AtNIP6;1* was not particularly differentially expressed in response to B-deficiency in any of the accessions, with no genotypes exhibiting greater than 2-fold up- or downregulation. On average across all analysed genotypes, there was a 12% reduction in expression of *AtNIP6;1* in B-deficient compared to B-sufficient conditions. There was also no difference in expression response between B-efficient or B-inefficient genotypes (*p* = 0.99). *AtBOR1* was on average upregulated 1.38-fold under B-deficient conditions, although expression was more varied between accessions than *AtNIP6;1*. Up- or downregulation varied 5.3-fold among B-efficient genotypes from 0.85 to 4.46, and 3.6-fold among B-inefficient genotypes from 0.72 to 2.55 (Fig. **5b**). Again, there was no difference in the *AtBOR1* expression response between B-efficient or B-inefficient genotypes (*p* = 0.98). No significant correlations were detected between expression of any of the three analysed B transporters and BEI_root_ or BEI_shoot_. Coupled with the lack of significant difference in expression between B-efficient and B-inefficient genotypes, this indicated that altered expression of B transporters is not the main driver of B-efficiency in *A. thaliana*. In addition, the root expression levels of the three analysed B transporters did not significantly correlate with shoot B concentration in either B-deficient or B-sufficient growth conditions, indicating that transporter-driven B uptake was already at a maximum for all assessed accessions.

### Boron transporter proteoforms do not differ between B-efficient and B-inefficient *A. thaliana* accessions

As the transcript abundance of the three B transporters AtNIP5;1, AtNIP6;1 and AtBOR1 did not correlate with shoot biomass accumulation under B-deficient growth conditions between selected accessions contrasting in B-deficiency tolerance, we tested for correlations between the amino acid sequences (proteoform layout) of these B transporters and B efficiency. The AtNIP5;1 amino acid sequences of the accessions investigated in this study were shared with that of Col-0 with the exception of Petergof (V_299_ to I_299_) and Zdr-1 (M_12_ to V_12_). The frequencies of these proteoforms among the 1135 Arabidopsis accessions of the analysis tool were 0.09 and 0.44%, respectively, indicating that these protein variants are rare. Moreover, they were not associated with B-inefficiency. The AtNIP6;1 variants V_259_ to I_259_ and G_193_ to D_193_ also exist among the accessions of the current study, which occur with a frequency of 24.05 and 28.9% across the set of 1135 accessions of the analysis tool. However, these variants occur both in accessions with a high (0.9 percentile: V_259_ to I_259_: Eden-1, Zü-1; G_193_ to D_193_: Yo-0) and a low (0.1 percentile: V_259_ to I_259_: Hovdala-2, War, Bay-0; G_193_ to D_193_: Van-0, Hau-0, Ull2-3, Kin-0) BEI_shoot_. Similarly, 37.27% of the 1135 Arabidopsis accessions carry a L_548_ to I_548_ exchange in AtBOR1. At least 48 accessions of the accessions in this study carry this allele, which also correlated with neither BEI_root_ nor with BEI_shoot_ and is present both in B-efficient and B-inefficient accessions. Additionally, the accessions Stw-0 and Wil-2 had a D_399_ to E_399_ alteration and Tamm-2 had a V_78_ to G_78_ variant in AtBOR1, both representing rare proteoforms.

### Boron-efficient accessions cluster phylogenetically and geographically

A phylogenetic clustering approach based on 214,051 SNPs was used to identify shared genetic bases and evolutionary origins of B-efficiency in *A. thaliana*. The full dendrogram is shown in Fig. **S6**. There was good clustering observed between genotypes of known similar origin, including the accessions N4, N7 and N13 from the Republic of Karelia, Russia, Ws, Ws.2 and Ws.3 from Wassilewskija, Russia, and Dra.0, Dra.2, Da.1..12 and DraIV6.35 from Czechia. We detected 11 distinct clades within the dendrogram (Fig. **7a**). A clear enrichment of B-efficient accessions was observed in clade 6 of the dendrogram, making up seven of the 14 accessions within this clade (Fig. **7a**). This included three out of the five accessions defined as B-efficient based on both BEI_shoot_ and BEI_root_ scores (Fig. **7b**). Five of the remaining seven accessions in clade 6 were also well within the 70^th^ BEI_shoot_ percentile with BEI_shoot_ scores upwards of 0.31, and were therefore also relatively B-efficient compared with the rest of the accessions. Interestingly, the two accessions Tamm-2 and Tamm-27, which had BEI_shoot_ scores of 0.11 and 0.07, respectively, and therefore were relatively B-deficiency sensitive accessions, also fell within this clade. Both of these accessions originate from Tammisari, Finland. The genotype Es-0, which we characterised as a B-efficient genotype with a BEI_shoot_ of 0.54, also originates from Finland and was sampled from only approximately 70 km away. It is likely that these accessions share a common ancestor but diverged in their B-efficiency more recently. Three other accessions from clade 6, Eden-1, Var2-1 and Ost-0, originated in Sweden and therefore relatively nearby to the previously mentioned accessions of this clade, further suggesting a shared recent ancestor for these accessions. Areas of Scandinavia were previously characterised as having B-deficient soils (Shorrocks, 1997), which may have provided a selection pressure for adapting to low B environments in these genotypes.

**Fig. 7.**
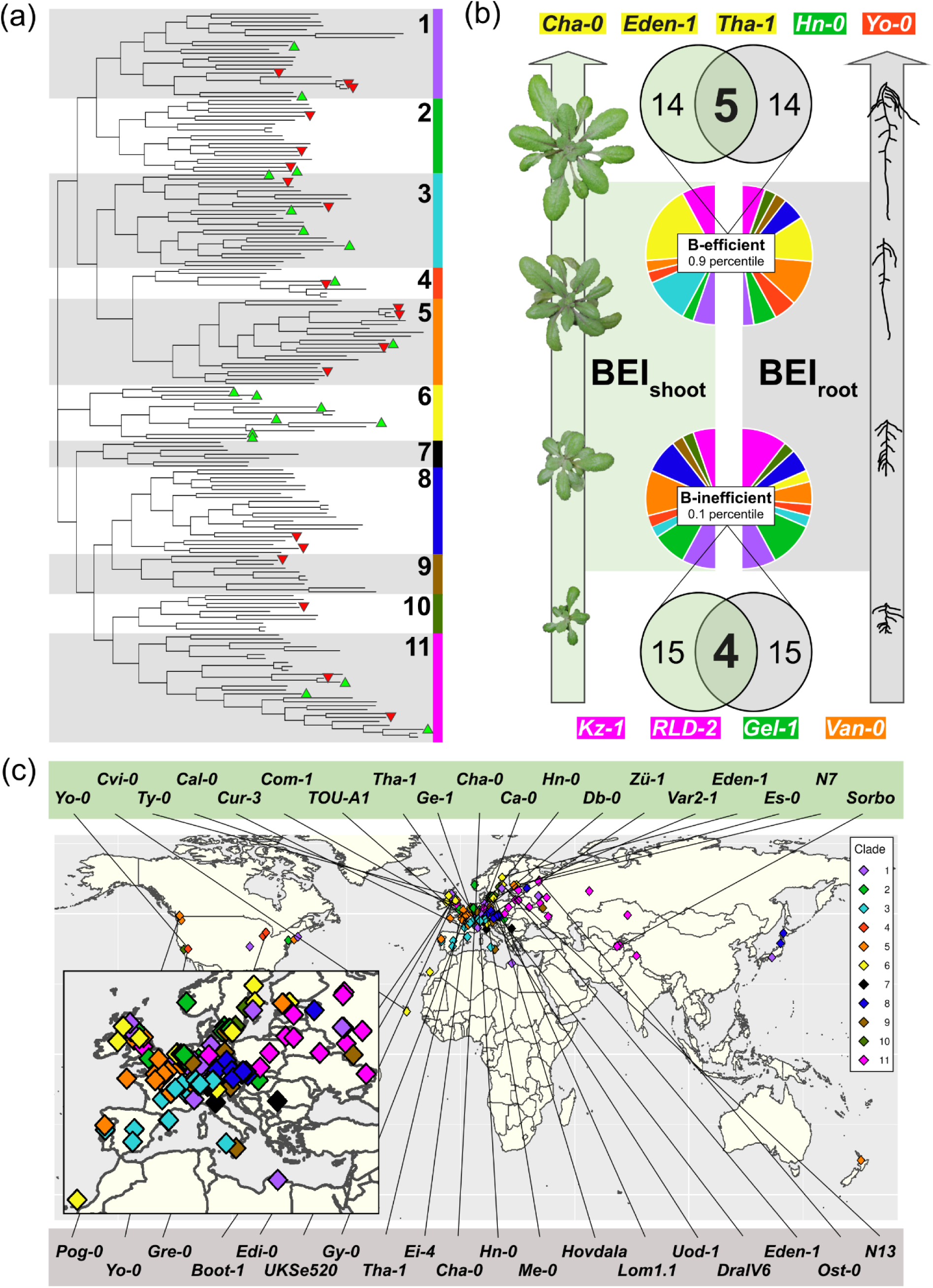
**(previous page).** Phylogeny and geography of B-efficiency. (a) Dendrogram showing phylogenetic relationships of the 185 *A. thaliana* accessions, with 11 clades manually assigned. Green triangles represent B-efficient accessions of the BEIshoot 90^th^ percentile (n = 19). Red triangles represent B-inefficient accessions of the BEIshoot 10^th^ percentile (n = 19). Full dendrogram with genotype names show in Fig **S6**. (b) Proportion of accessions of the BEIshoot and BEIroot 10^th^ and 90^th^ percentiles that exist within each phylogenetic clade. Colour-coding in pie charts represents phylogenetic clades. Sizes of slices of pie indicate more or less genotypes of the corresponding clade that are in either of the 10^th^ or 90^th^ percentiles. Accessions listed at the top or bottom of the panel are those that were ranked consistently B-efficient or B-inefficient, respectively, in both shoot and root datasets, colour-coded by phylogenetic clade. (c) Location of sampling point of all 185 accessions colour-coded by phylogenetic clade. Accessions of the BEIshoot (top, green background) or BEIroot (bottom, grey background) 90^th^percentile indicated.

Clade 3 contains four further accessions characterised here as B-efficient (CUR-3, TOU-A1-116, Zü-1, Ge-1), as well as two accessions defined as B-intermediate efficient accessions (Bs-1, Co). Five out of six of these originated from either France or Switzerland, indicating once more B-efficiency acquired through a shared common ancestor. An additional two and one accessions of this clade also originated from France or Switzerland, respectively, and five from Germany close to the French and/or Swiss border, indicating a shared genetic origin prior to emergence of B-efficiency in this clade. No patterns of a shared genetic basis of B-inefficiency were immediately obvious from the phylogeny of the 185 accessions analysed here. However, two of the four accessions that had consistently low BEI scores in both shoots and roots, Kz-1 and RLD-2, both clustered in clade 11 (Fig. **7b**). Kz-1 originates from Kazakhstan and RLD-2 from Iran. These are regions that were not very well represented in our accessions set, but the poor performance of these genotypes in B-deficient conditions may indicate a poor selection pressure for B-efficiency.

The origin or sampling site of all accessions with co-ordinate data available (183 out of 185 accessions) is shown on a world map in Fig. **7c**, colour-coded by the phylogenetic clades indicated in Fig. **7a**. A clustering of accessions from clade 6 can be seen around Nordic countries. This includes the B-efficient lines from Sweden and Finland mentioned above, as well as Ty-0 from the United Kingdom and Tha-1 from the Netherlands. Accession Cvi-0 from Cabo Verde also ranked as a B-efficient accession based on BEI_shoot_ and clustered into this clade. In general, other phylogenetic clades clustered into distinct geographic regions, including clade 1 which included accessions mostly from Eastern Europe and Western Asia, clade 3 which included accessions mostly from Central, Southern and Western Europe, and clade 8 which included accessions mostly from Central Europe (Fig. **7c**).

Nineteen of the 185 accessions were sampled from altitudes above 500 m, of which 10 are in the 70^th^ BEI_shoot_ percentile, and six were sampled from above 1000 m, of which three were classified as either B-efficient or intermediate-B efficient in this study. There may therefore by a tendency for higher altitude accessions to be more B-efficient, and a significant trend between BEI_shoot_ and altitude was observed (Pearson’s correlation coefficient = 0.25, *p* < 0.001).

### Phenotypic response profiling revealed 12 distinct adaptive root and shoot performance patterns under B-deficiency

A subset of 38 trait ratios between B-deficient and B-sufficient growth conditions originating from automated phenotyping (n = 30), RGR and DW determination (n = 3) from the soil-substrate experiment, and from the Petri-dish experiment (n = 5; Fig. **S7**), were used to cluster genotypes into 12 *k*-clusters. The aim was to identify patterns of responses to B-deficiency among the 185 accessions, and to identify which traits most associated with high or low growth performance in low B environments. Cluster 6 contains 28 accessions that together had the highest average DW in the automated phenotyping experiment compared to other clusters (Fig. **8**). Eleven of the 19 B-efficient genotypes in the 90^th^ percentile of BEI_shoot_ were also grouped into cluster 6. Accessions within this cluster had relatively high values across all traits measured, with the exception of the colour traits brown:green and yellow:green pixel ratios, and lateral root number. These colour traits are negatively associated with biomass accumulation in B-deficient compared to B-sufficient conditions (Fig. **S8**), and were generally highest in *k*-clusters with low average DW and PLA (Fig. **8**). The centered and scaled average (*k*-mean) LRN of accessions within cluster 6 is close to zero, and as such, this trait is not associated with the high B efficiency among accessions of this cluster. However, all other root traits had relatively high *k*-means in this cluster, most notably ALRL and TLRL. Both ALRL and TLRL are strongly correlated across the accessions, and both are generally correlated with shoot biomass accumulation traits (Fig. **S8**). The clusters 3 and 4 which also exhibited a high *k*-mean TLRL also had a higher than average ratio of shoot DW between B-deficient and B-efficient accessions. However, different B-efficiency mechanisms appear to be present in other *k*-clusters. Cluster 1 (n = 21) is characterised by a relatively good shoot performance under B-deficient conditions but has only average TLRL and ALRL and a high average PRL and LRN, indicating that these traits may contribute to B-efficiency in some accessions.

**Fig. 8.**
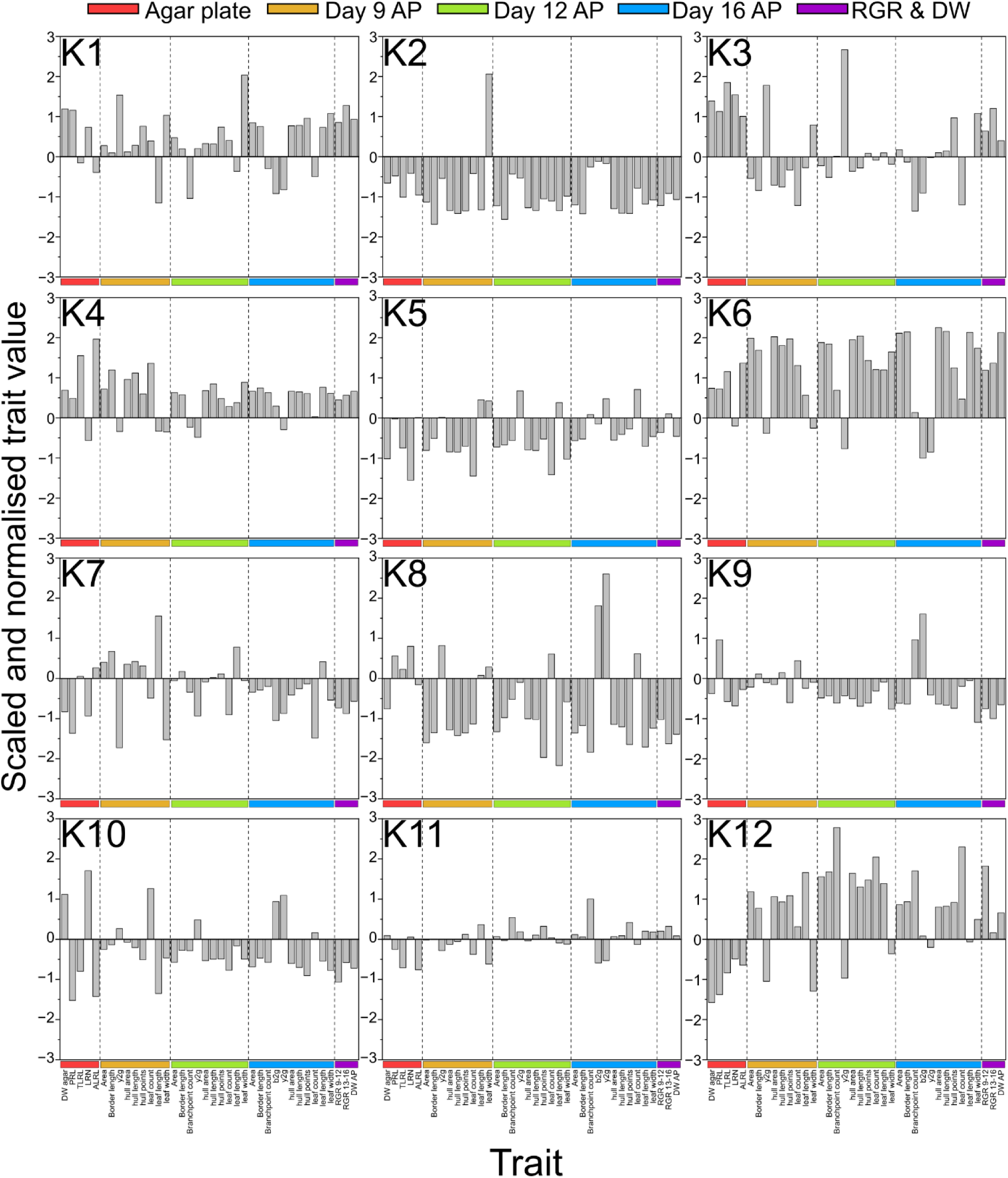
Centered and scaled trait ratios between B-deficient and B-sufficient growth conditions of the 38 traits included in *k*-means clustering. The trait ratios of the 12 *k-*clusters are shown separately. Centering was performed by subtracting the trait ratio mean from each accession ratio. Scaling was performed by dividing this value by the trait ratio standard deviation.

Cluster 12 also maintained a relatively good shoot performance under B-deficiency, but with generally very poor root performance. Only five accessions were grouped into this cluster (Com-1, Kondara, Ler-1, Oy-0 and Tsu-0). Com-1 and Kondara have relatively high BEI_shoot_ scores. However, none of these accessions clustered into the same phylogenetic group, so a shared genetic basis of B-efficiency in these genotypes is unlikely. Overall, maintaining a large root system under B-deficient conditions and in particular producing longer but not more lateral roots, appears to be an efficient strategy to maintain shoot productivity in such conditions.

## Discussion

### Boron efficiency is rare amongst *A. thaliana* accessions

Of the 185 accessions included in the present study, we classified the 19 in the 90^th^ BEI_shoot_ percentile as B-efficient. However, of these, only nine maintained both PLA at day 16 and shoot DW to at least 80% of their level under B-sufficient conditions when exposed to B-deficiency. Only three accessions, Var2-1, Hn-0 and Tha-1 maintained both traits above 90% of their respective levels under B-sufficient conditions when exposed to B-deficiency. We therefore consider B-efficiency to be rare in *A. thaliana*. This is perhaps not so surprising, since most natural soil in the world is not B-limiting and if B-deficiency occurs, this is generally sporadic and temporally-confined (Brdar-Jokanović, 2019; Landi et al., 2019; Shorrocks, 1997; Gupta et al., 1985). The three above-mentioned accessions originated from Sweden, Germany and Netherlands, respectively. Parts of northern Europe, particularly Scandinavia, have been characterised as having low soil B availability (Shorrocks, 1997), and so it is also not particularly surprising that B-efficient accessions were found here. Var2-1 and Hn-0 are two out of the three accessions that maintained at least as high shoot DW in B-deficient compared to B-sufficient conditions. The third accession is Yo-0 which was sampled from Yosemite National Park in California, USA at an altitude of 1,400 to 1,500 m. High altitude has been linked to reduced B concentration and availability in India (Kumar et al., 2019; Pradhan et al., 2003). It is therefore feasible that the region from which Yo-0 was sampled had low soil B availability and therefore created a selection pressure for B-efficiency.

### Lateral roots contribute most to B-efficiency among root traits

In contrast to the soil-substrate growth system, 36 accessions maintained at least as much shoot biomass in B-deficient compared to B-sufficient conditions in the agar-based Petri-dish cultivation system, and a further 61 maintained shoot biomass above 80% of their respective levels under B-sufficient conditions. This indicates that B was not yet a major factor limiting seedling shoot growth in B-deficient conditions in this system. The Petri-dish cultivation system was optimised for growing and quantifying root system architecture. None of the included accessions maintained as much as 80% of their PRL in B-deficient compared to B-sufficient conditions, and only nine maintained above 70% PRL (Fig **6**). Boron-limitation is known to have a detrimental effect on the root system architecture and root functioning of plants (Liu et al., 2021). In a previous study, the length of *A. thaliana var.* Col-0 primary roots were up to 39% shorter in an environment with no added B compared to control conditions, whilst lateral root density and growth slightly increased (Gruber et al., 2013). Primary root length was also the most negatively impacted root trait measured in the present study, indicating either a requirement for B to elongate the primary root and to sustain mitotic activity of the root meristem or a response to low B that favours investment in other plant organs. In support of the latter, 16 accessions increased their TLRL, and 38 increased their ALRL under B-deficient compared to B-sufficient conditions. In many soils, more B is found in the topsoil where it is associated with soil organic matter (Gupta et al., 1985). Increasing lateral root length may therefore represent a favourable strategy to acquire more B where soil concentrations are low, as also indicated by the *k*-clustering analysis (Fig. 8). However, B is also relatively mobile in soil and can easily leach to lower soil layers especially after strong rainfall (Brdar-Jokanović, 2019; Shorrocks, 1997). In such situations, growing longer roots may be a favourable strategy, although this may be more effective in larger and deeper rooting plants than *A. thaliana*. Interestingly, the accession Var2-1 which had the highest BEI_shoot_ displayed the second longest PRL under B-deficient growth conditions but the second shortest TLRL among accessions here. This suggests that in this accession, the lateral roots did not significantly contribute to B-efficiency, therefore revealing variation in B-efficiency strategies between accessions.

Aside from the possibility that the decrease in primary root length represents an adaptive response to B-deficiency to favour resource allocation to lateral roots, it may also be attributed to reduced cell wall integrity caused by a lack of B-bridging between rhamnogalacturanon-II monomers. Indeed, it has been shown that B-deficiency leads to a signalling pathway involving ethylene, auxin, and ROS to restrict root cell elongation in a manner that resembles cell wall stress responses (Camacho-Cristóbal et al., 2015). Whilst some accessions here did exhibit increased lateral root length under B-deficiency, the length of the overall root system was decreased in all but two accessions (Ei-4 and DraIV6-35), with 180 out of 185 accessions suffering more than a 20% decrease in total root system length. Therefore, a reduced cell wall integrity leading to reduced root cell elongation and/or reduced mitotic activity of the root meristem remains a reasonable theory for the root response to B-deficiency observed in most accessions here.

### In most accessions, B-efficiency is neither associated with increased B transporter expression nor B transporter sequence variation

Shoot B concentrations of the 22 analysed B-efficient accessions were on average 41% higher than the 21 analysed B-inefficient accessions under B-deficient growth conditions. These B-efficient accessions also accumulated 4.8 times more shoot DW under B-deficient conditions than B-inefficient accessions, and so accumulated 6.5 times more shoot B per plant on average. Remarkably, no differences between expression of the B transporters *AtNIP5;1*, *AtNIP6;1* or *AtBOR1* were observed between B-efficient and B-inefficient accessions. Three remaining possibilities exist for how the B-efficient accessions were able to accumulate so much more B in shoot tissue: a bigger root system, more effective transport protein alleles, or increased mass-flow of B towards root surfaces for increased passive and active B uptake. Increased B accumulation early in development when more B is available in the soil is unlikely to be an effective growth strategy, since B is generally immobile in most plant species and cannot readily be retranslocated (Brown & Shelp, 1997). Therefore, an early depletion of soil B would likely result in more severe B-deficiency symptoms being experienced later. Of the 22 B-efficient accessions and 21 B-inefficient accessions analysed for expression of *AtNIP5;1*, *AtNIP6;1* or *AtBOR1*, the B-efficient accessions produced 25% longer whole root systems in the agar-based Petri-dish cultivation system than the B-inefficient accessions, on average. The gene expression measurements considered the relative expression of the target B transporters per unit of RNA. A larger root system means more cells and therefore more RNA, which implies increased abundance of B transport proteins in accessions with larger root systems even when the relative expression between accessions is the same. However, it is unlikely that the average of 25% longer root length in the B-efficient accessions alone explains the 6.5 times greater B accumulation in these accessions. More effective transport protein alleles are also unlikely to explain the difference in B accumulation, since none of the few amino acid substitutions we observed between accessions correlated with B-efficiency. From the data we collected here, it is impossible to say if variation in mass flow of B towards root surfaces, for example through increased transpiration (Oyewole et al., 2014), existed between B-efficient and B-inefficient accessions. Further work will be required to determine whether such differences exist between *A. thaliana* accessions and if these contribute to increased B uptake, and in addition, whether B-efficient accessions would benefit from a temporal increase in B availability more than B-inefficient accessions. However, even B-efficient accessions suffered an 84% drop in shoot B concentration under B-deficiency, compared to an 87% drop in B-inefficient accessions. Therefore, additional B-efficiency mechanisms beside effective B uptake must control the B-deficiency tolerance of these accessions.

### Boron efficiency evolved independently in different regions

Of the 185 accessions included here, 153 are described as having been sampled from different countries in Europe. It is therefore perhaps unsurprising that of the 30 accessions characterised as B-efficient or B-intermediate efficient based on BEI_shoot_, 24 originated from Europe. Of the remaining efficient accessions, three originated from Asia (Sorbo and Kondara, Tajikistan; Kas-2, India), one from Africa (Cvi-0, Cabo Verde), one from North America (Yo-0, USA), and one was not defined (S96). Similarly, 16 out of 19 accessions in the 90^th^ percentile for BEI_root_ originated from Europe, with the remaining three from the USA (Yo-0, Gre-0) or Canada (Pog-0). Despite the relatively confined geographic spread, it is clear that not all of the identified B-efficient accessions originated from a single B-efficient ancestor. Hotspots for B-efficiency certainly exist, most notably in Scandinavian countries in which the B-efficient accessions Var2-1, Es-0 and Eden-1 cluster together both genetically (clade 6, Fig. **7a**) and geographically (Fig. **7c**). Three of the remaining four B-efficient accessions of clade 6 also originated from Europe, and one from Cabo Verde. Phylogenetic clade 3 contains a further six European accessions defined here as either B-efficient or B-intermediate efficient, originating from Switzerland (3), France (2) or Portugal (1). However, the different phylogenetic grouping suggests that these accessions originated from a different B-efficient ancestor. Clade 11 accessions were more diverse in origin, with B-efficient or B-intermediate efficient accessions in this clade originating from Tajikistan (2), the United Kingdom (1), Lithuania (1), Russia (1) or India (1). Longer distances between some of the accessions may have been human transport influenced. Despite these accessions all having relatively high BEI_shoot_ values, their performance for other traits varied between the phylogenetic groups. Boron-efficient or B-intermediate efficient accessions from clade 11 had 45% and 57% more shoot DW under B-sufficient conditions than B-efficient or B-intermediate efficient accessions from clades 3 and 6, respectively, and 50 and 37% more shoot DW under B-deficient conditions. This appears to be controlled by a longer TLRL in such accessions from this clade than from the accessions from both of the other clades. Meanwhile, B-efficient or B-intermediate efficient accessions from clade 6 had the longest PRL, as well as the highest shoot B concentration under B-deficient growth conditions. Interestingly, both clade 3 and clade 11 also contain two and three accessions, respectively, that were in only the 10^th^ BEI_shoot_ percentile, and that were therefore among the most B-deficiency sensitive accessions identified. Clade 6 also contains the accessions Tamm-2 and Tamm-27 that were generally much more B-deficiency sensitive than the other accessions in the clade. Therefore, B-efficiency appears to have evolved within each clade separately, representing at least three independent adaptations across the described geographic areas. The remaining 11 B-efficient and B-intermediate efficient accessions based on BEI_shoot_ were scattered throughout seven different phylogenetic clades and seven countries and may represent further independent adaptations to low B-availability.

Aside from northern Europe, Shorrocks (1997) also identified localised regions of North America, South America, Africa and eastern Asia that had B-deficient soils. Co-localising with two of the B-deficient regions in North America are the accessions Pog-0 and Gre-0 which were both in the 90^th^ percentile for BEI_root_. However, neither of these accessions had particularly strong shoot performance under B-deficiency. In addition, other accessions in the same phylogenetic clades localised very closely to these two accessions but were considered B-efficient neither based on BEI_shoot_ or BEI_root_. Therefore, we cannot comment overall as to whether genotypes from these regions are generally B-efficient. No B-efficient accessions were identified from the other B-deficient regions from Shorrocks (1997) since no accessions from these regions were included in the analysis here. Additional work with accessions from these regions will be required to discover whether B-efficiency evolved independently there as well.

Follow up studies have been initiated aimed at identifying genetic components modulating *A. thaliana* root and shoot growth under B deficiency. Approaches being implemented include genome-wide association mapping and QTL mapping taking advantage of a recombinant inbred line population derived from a cross between two very closely related clade 11 accessions, namely Rld-2 a B-efficient accession, and Cal-0, a B-inefficient accession.

## Conclusion

Our comprehensive analyses provide quantitative evidence for phenotypic and molecular plasticity in root and shoot responses to B-deficiency within 185 *A. thaliana* accessions. Whilst B-efficiency in *A. thaliana* is rare, we successfully identified several B-deficiency tolerant accessions that appear to have evolved distinct and separate B-efficiency mechanisms. Co-localisation of many B-efficient accessions with regions previously characterised as B-deficient indicates that B-efficiency evolved as an adaptation to such environments. Overall, our analysis indicates root system architecture and in particular increased lateral root length but not number contributes to B-efficiency in many but not all B-deficiency tolerant accessions. Elemental analysis and gene expression analyses revealed slight differences in B concentration between B-efficient and B-inefficient accessions but no differences in expression of major B transporters. This indicates a low B requirement of B-efficient accessions and therefore an increased B-utilisation efficiency. The existence of close genetic relationships between B-efficient and B-inefficient accessions, such as Rld-2 and Cal-0 which clustered adjacently to each other in the phylogenetic tree, but fell into the 10^th^ and 90^th^ BEI_shoot_ percentiles, respectively, suggests that small genetic changes can have a large effect on B-efficiency. The large volumes of shoot imaging and root phenotyping data generated here are considerable and will be useful for subsequent studies aiming to improve the B-resource efficiency and B-status of crop plants grown under suboptimal B conditions. The challenge now will be to combine datasets and genomics resources to pinpoint genetic variation and molecular and metabolic mechanisms controlling variation in B-efficiency.

## Acknowledgments

We gratefully acknowledge the specified funding sources. We acknowledge Dr. Yudelsy Tandron Moya (IPK) for excellent technical assistance in the ICP-MS analysis, Jacqueline Fuge (IPK) for RT-qPCR analysis and Ingo Mücke (IPK) for his support in management of the “IPK automated plant phenotyping system for small plants” system operations. We also thank Annett Bieber, Jacqueline Fuge, Kristin Vorpahl, Benjamin Pommerrenig, Arvid Diehn, Nicola Mancioppi, Agostino de Matteis, Gunda Wehrstedt, and Beatrice Knuepfer for their excellent assistance in the growth of plants and data processing. This work was supported by the Emmy Noether grant 1668/1-1 and 1668/1-2 from the Deutsche Forschungsgemeinschaft (to G.P.B.).

## Author contributions

GPB conceived the study. GPB, MDB and TDA wrote the first paper draft. All authors prepared, read, and approved the final manuscript. The B-deficiency tolerance screens in *in vitro* and *in terra* growth conditions and related experiments were conducted mainly by MDB and GPB but with the help of other authors. qPCR analysis and element analysis preparation were performed by MDB. Proteoform analysis was conducted by GPB and SK. Phylogenetic and bioinformatic analysis was performed by TDA. Work related to the automated imaging/phenotyping and subsequent analyses have been performed by AJ, MDB, TDA, TA, and GPB. GPB, MDB, NvW, TA, and AJ analyzed and interpreted the data.

## Supplementary Table

**Table S1.**
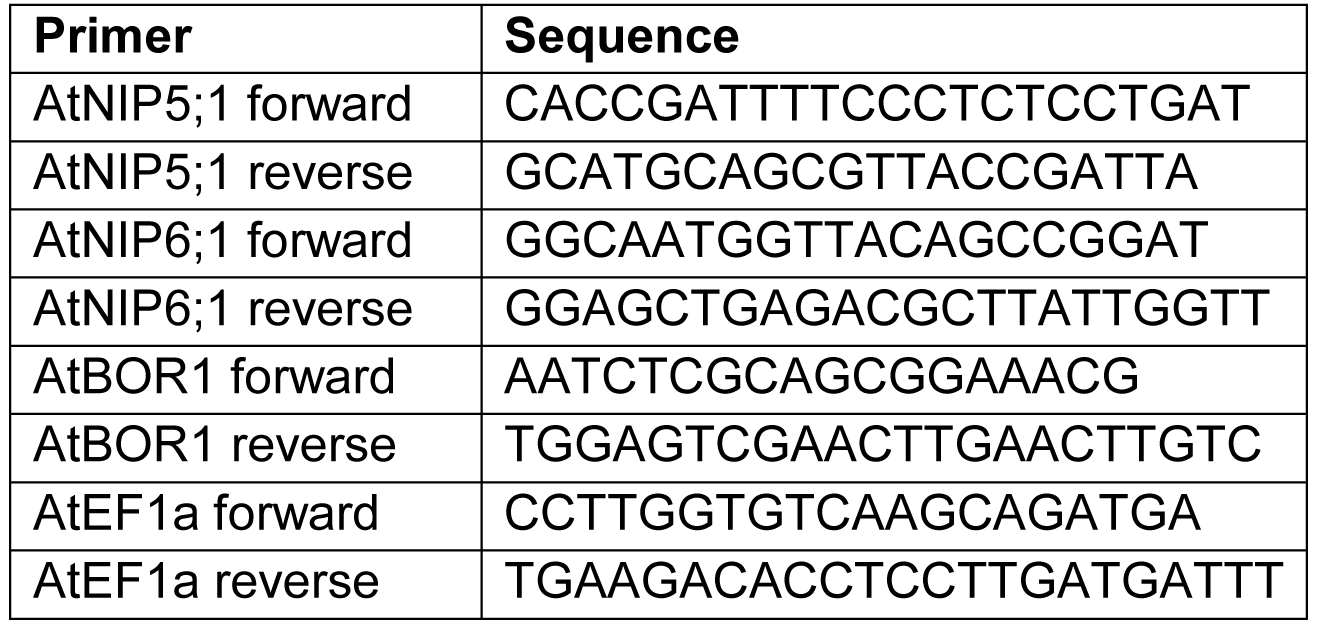
Primer sequences for RT-qPCR amplification of boron transporter encoding genes.

## Supporting Information Captions

**Fig. S1.** Correlations between area ratio between B-deficient and B-sufficient growth conditions on different imaging days, between visible light and fluorescence-based imaging, and between area at day 16 and dry weight (DW) at experiment end.

**Fig. S2.** Range in photosynthetic operating efficiency of each accession in both B-deficient (-) and B sufficient (+) growth conditions (a), and the genotypic ratio in performance between growth conditions (b). Boxes in (a) represent the 25^th^ to 75^th^ percentiles and whiskers represent the range within 1.5 times the interquartile range (IQR) with any outliers shown.

**Fig. S3.** Concentrations of plant macro-nutrients of 43 selected accessions considered B-efficient (22) or B-inefficient (21) in B-sufficient (PB) or B-deficient (MB) growth conditions. Boxes represent the 25^th^ to 75^th^ percentiles and whiskers represent the range within 1.5 times the interquartile range (IQR) with any outliers shown.

**Fig. S4.** Concentrations of plant micro-nutrients of 43 selected accessions considered B-efficient (22) or B-inefficient (21) in B-sufficient (PB) or B-deficient (MB) growth conditions. Boxes represent the 25^th^ to 75^th^ percentiles and whiskers represent the range within 1.5 times the interquartile range (IQR) with any outliers shown.

**Fig. S5.** Variation in relative (rel.) B transporter expression of the 43 selected accessions in B-sufficient (grey bars) or B-deficient (white bars) growth conditions. Colours by accession name represent phylogenetic clade as shown in Fig. 7.

**Fig. S6.** Full dendrogram showing phylogenetic relationship of the 185 *A. thaliana* accessions. Numbers in red at junctions refer to bootstrap scores.

**Fig. S7.** Pipeline to reduce traits measured across experiments to 38 informative traits for *k*-means analysis.

**Fig. S8.** Correlations table for all 38 traits included in *k*-means analysis.

