## Supplementary figures and images for "Natural variation in the *Arabidopsis thaliana* root and shoot response to boron deficiency reveals sensitive and responsive phenes and phylogenetic and geographic clustering of boron efficiency adaptations"

### Fig. S1

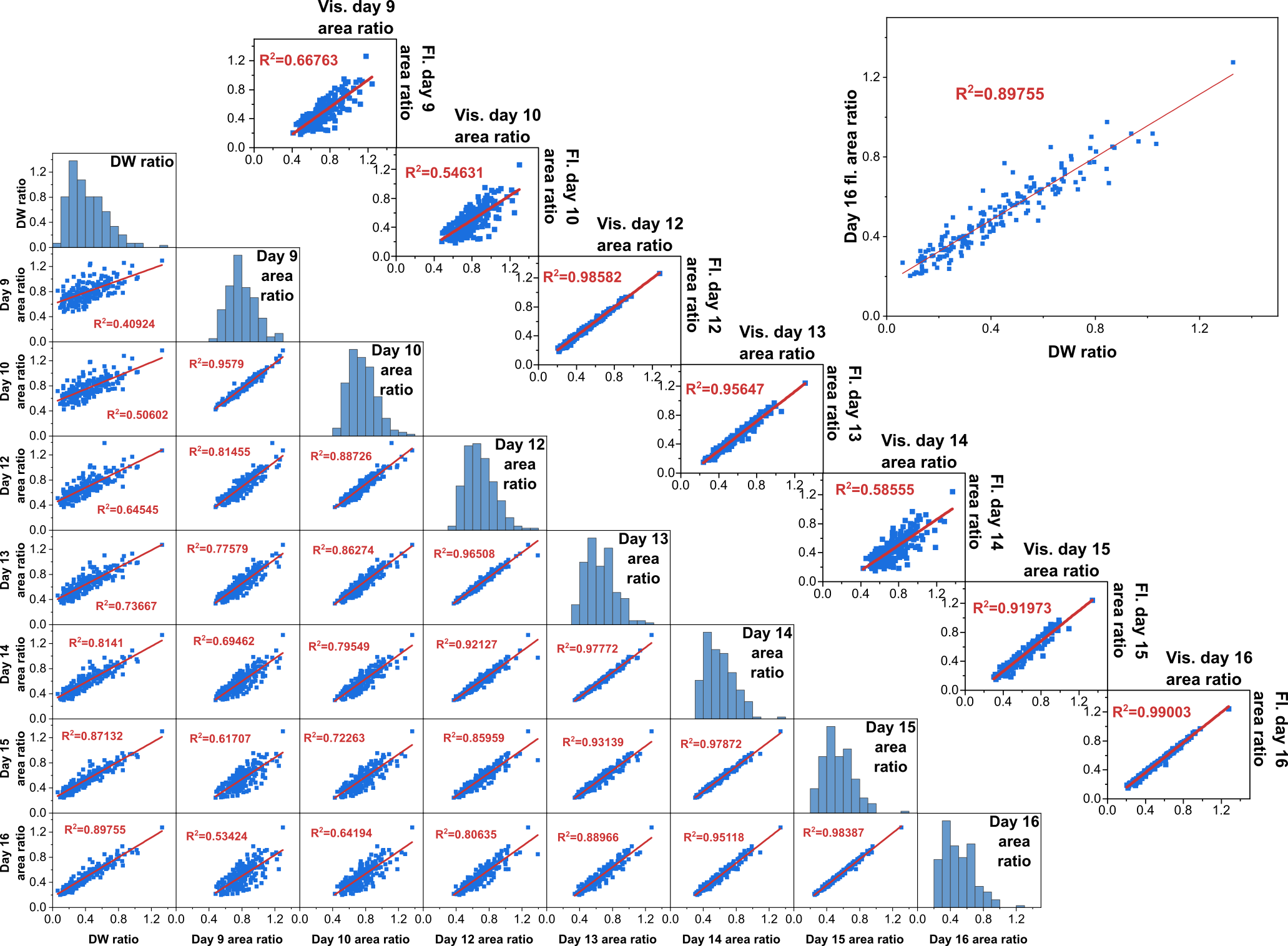

### Fig. S2

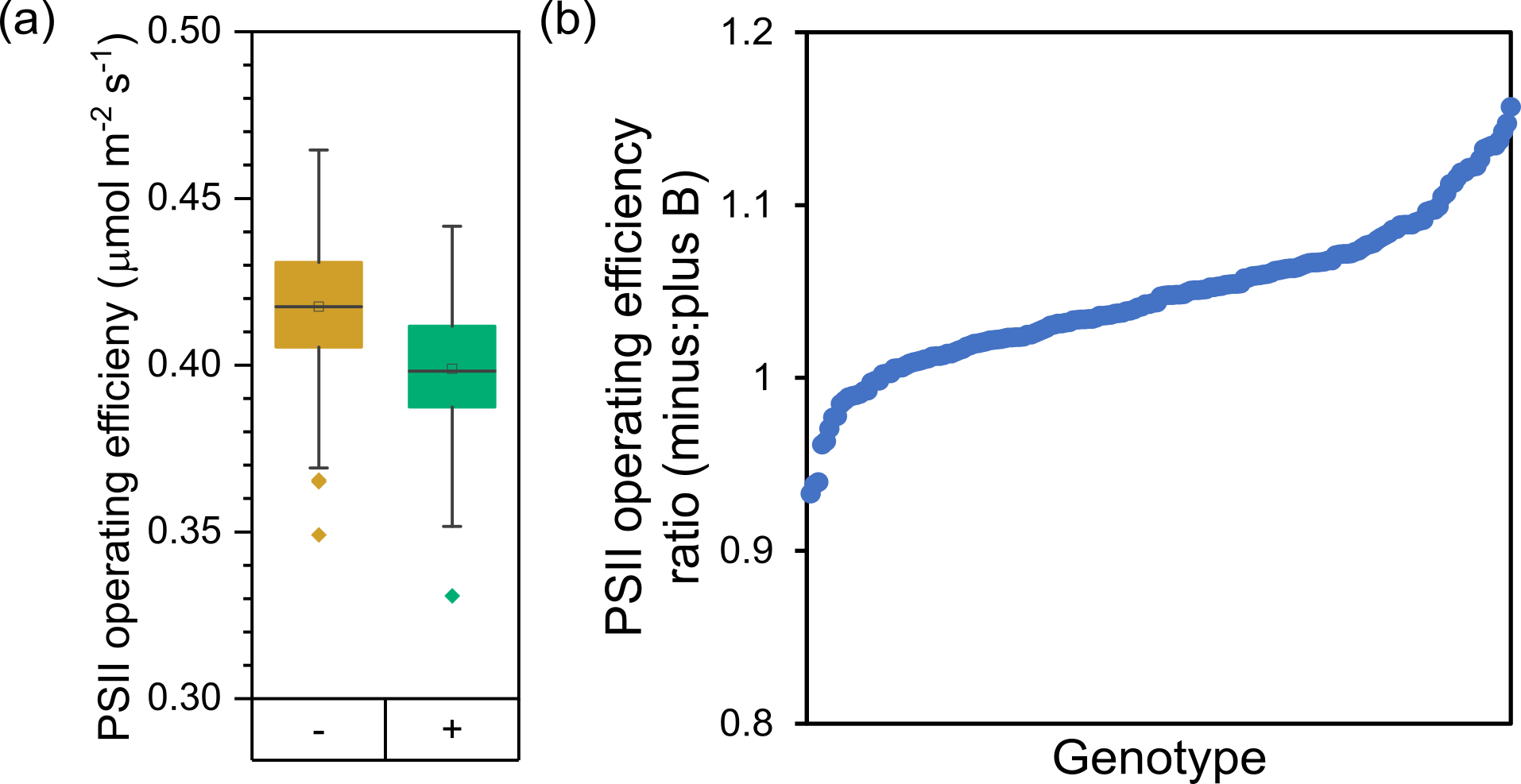

### Fig. S3

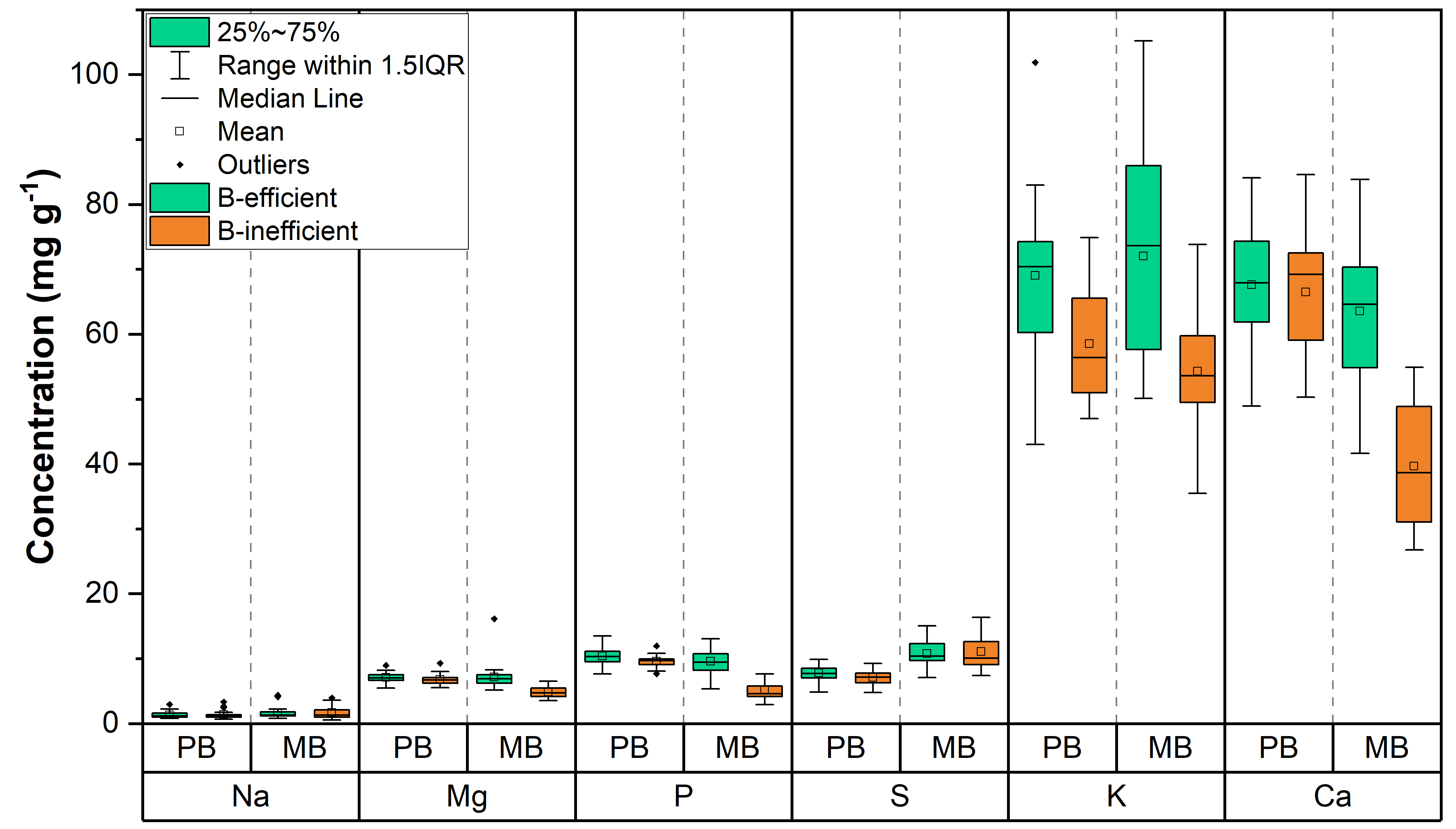

### Fig. S4

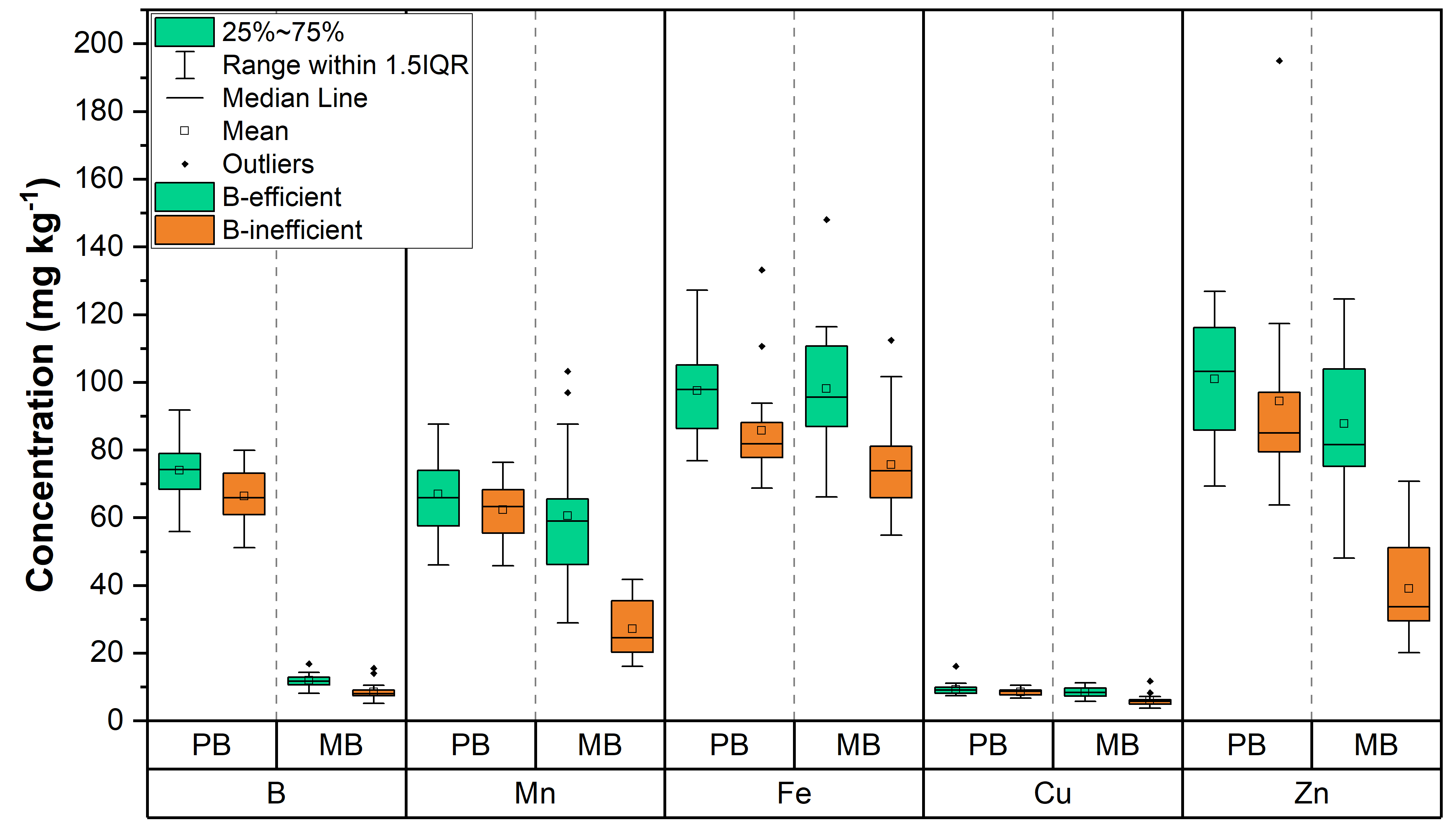

### Fig. S5

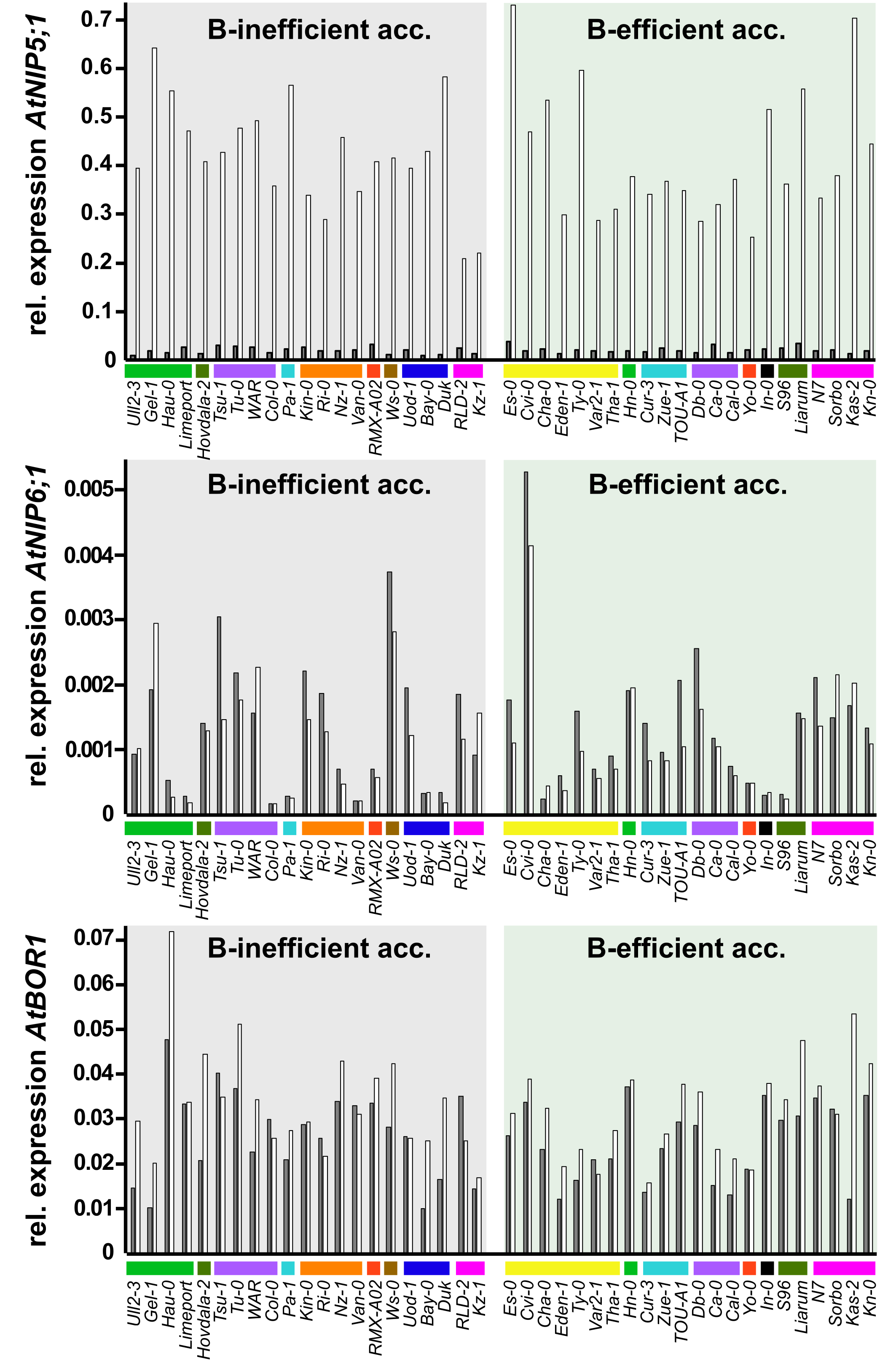

### Fig. S6

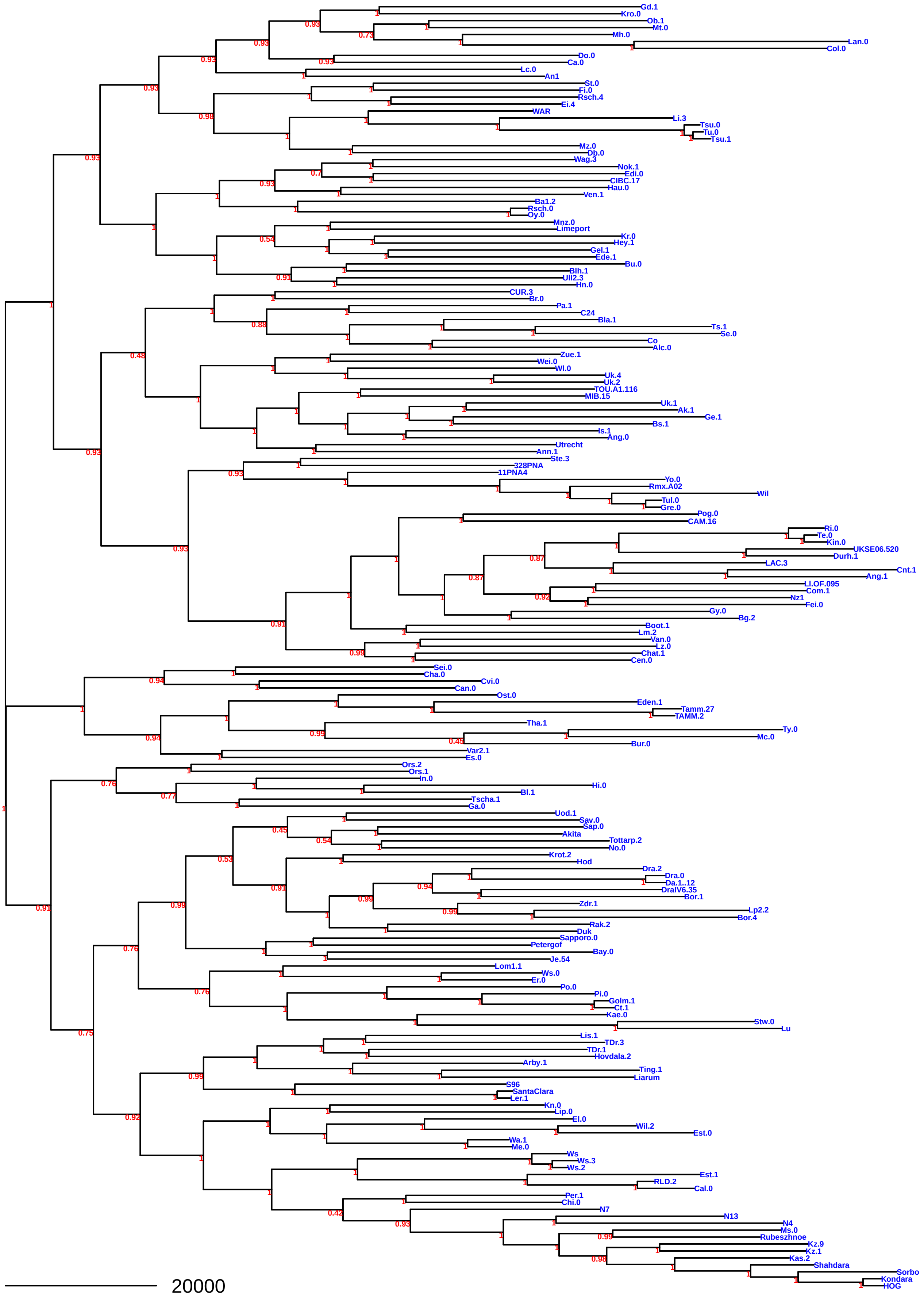

### Fig. S7

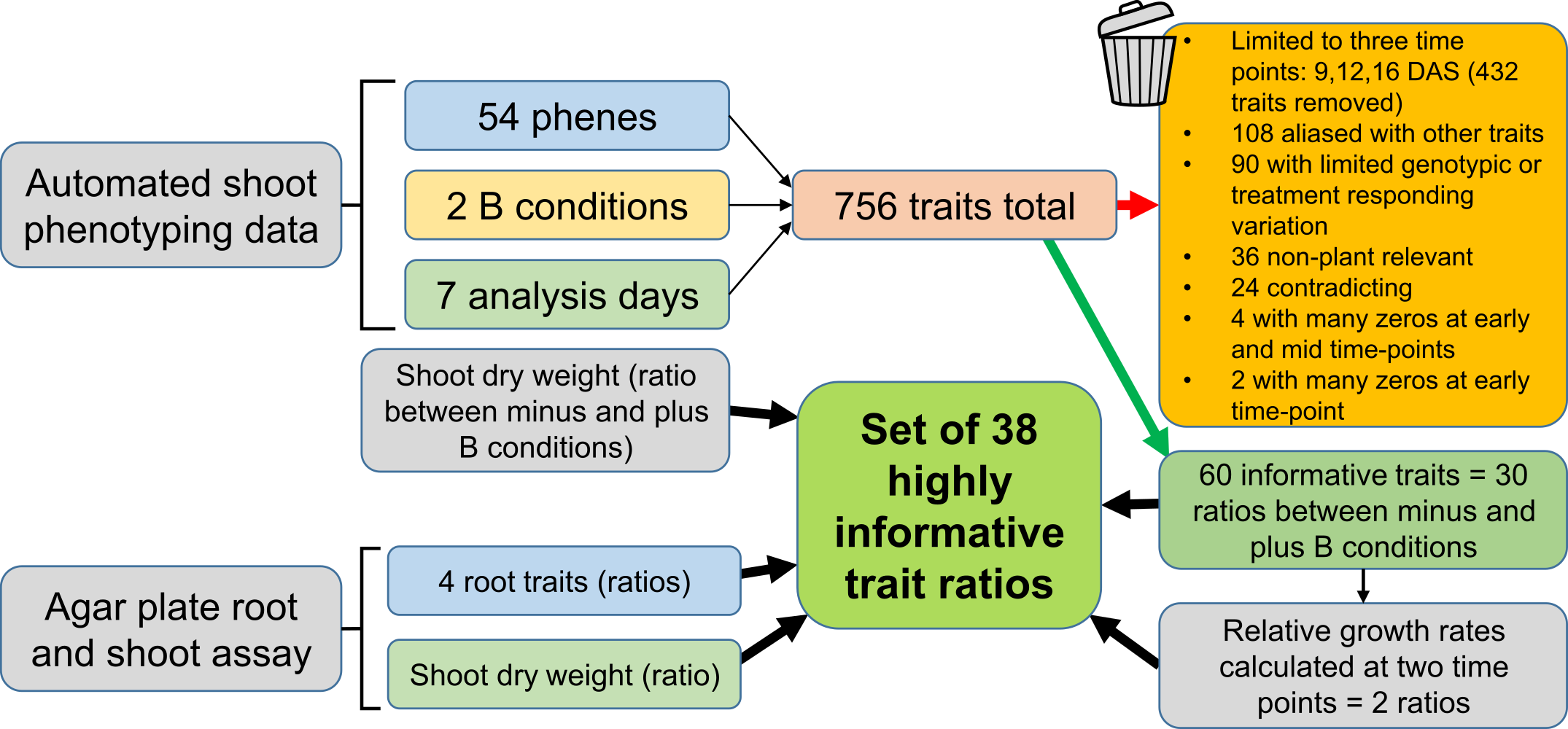

### Fig. S8

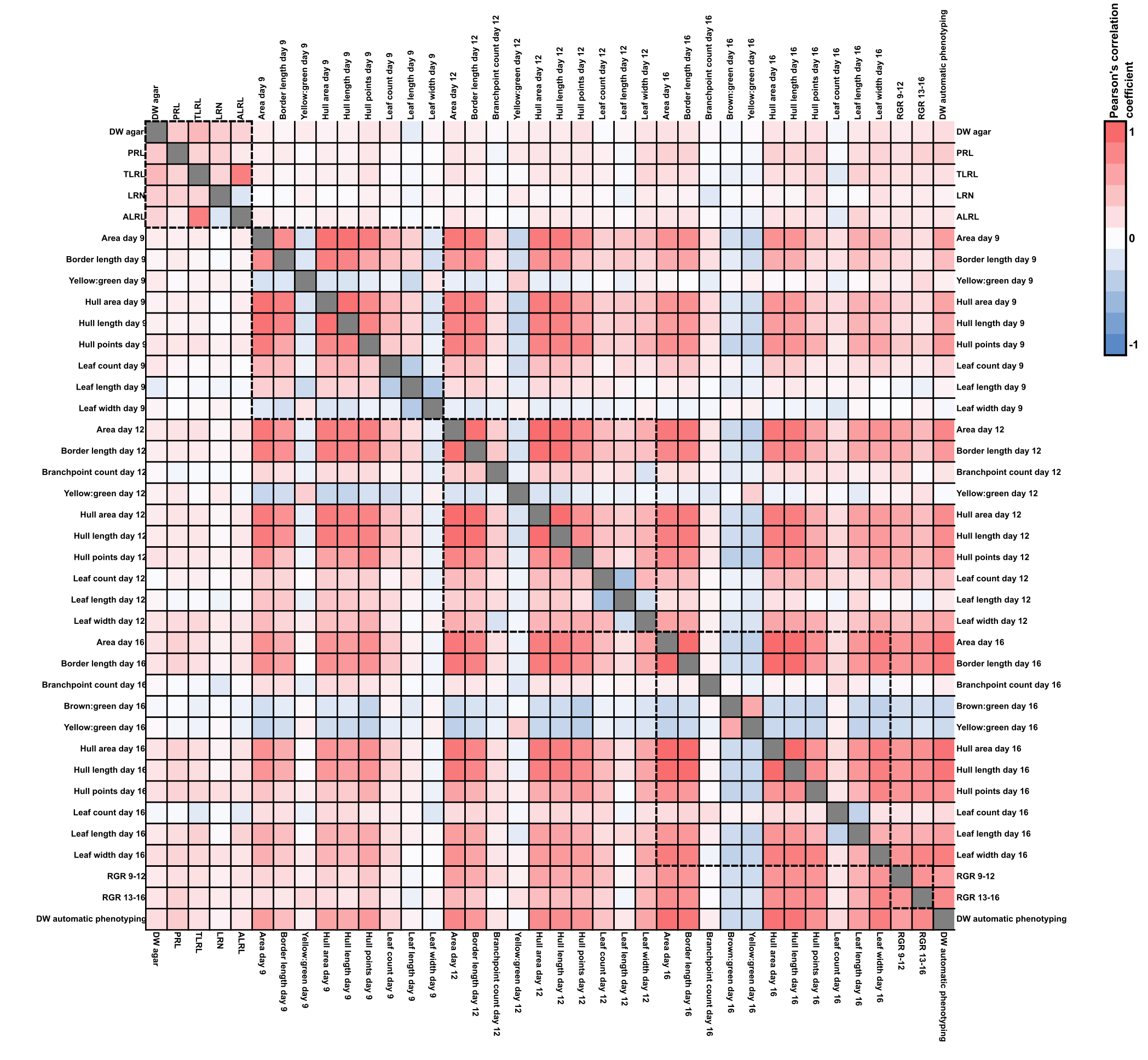
